# Distinct intersecting pathways link homolog pairing to initiation of meiotic chromosome synapsis

**DOI:** 10.1101/2024.03.11.584447

**Authors:** Jitka Blazickova, Shalini Trivedi, Richard Bowman, Sowmya Sivakumar Geetha, Silma Subah, Sarit Smolikove, Verena Jantsch, Monique Zetka, Nicola Silva

## Abstract

Faithful meiotic segregation requires pairwise alignment of the homologous chromosomes and Synaptonemal Complex assembly (SC) at their interface. Here, we investigate on new factors that promote and coordinate these events during *C. elegans* meiosis. We identify BRA-2 (<u>B</u>MP <u>R</u>eceptor <u>A</u>ssociated family member <u>2)</u> as an interactor of HIM-17, previously shown to promote double-strand break formation. We found that loss of *bra-2* specifically impairs synapsis licensing without affecting homologs recognition, SC maintenance or chromosome movement. Double mutant analysis revealed a previously unrecognized role for HIM-17 in promoting homolog pairing under dysfunctional SC assembly, without perturbing nuclear envelope recruitment of factors required for chromosome movement. We provide evidence that *bra-2* and *him-17* act in distinct pathways, exerting partially redundant functions in SC licensing, as well as separable roles in regulating homologs pairing. Altogether, our findings unveil novel mechanisms that ensure stabilization of homologous chromosome interaction via SC licensing upon homology assessment.

## INTRODUCTION

Formation of euploid gametes depends on the faithful execution of several disparate tasks during meiosis, all meticulously coordinated to ensure correct chromosome segregation into the daughter cells. Diploid genomes are composed of two homologous chromosomes, the paternal and the maternal copy, which must recognize each other (pairing) and recombine to achieve the formation of physical tethers called crossovers (COs). COs are essential to achieve successful segregation of the homologous chromosomes into the gametes^1,2^.

Pairwise chromosome alignment depends on microtubule-driven forces transmitted from the cytosol to the nucleus through the SUN-KASH protein bridge^3–6^. Once the chromosomes are connected to this protein module spanning the nuclear membranes, active movement along the nuclear envelope is triggered, enabling the chromosomes to travel within the nucleus until the synaptonemal complex (SC) has been established between homologous chromosome pairs. Therefore, chromosome movement is essential for pairing and effective homologous synapsis, as well as to resolve chromosome interlocks^3,7–9^.

In *Caenorhabditis elegans*, active chromosome motion occurs at meiotic entry, defining a specific region of the germ line called “transition zone” (TZ) that loosely corresponds to leptotene/ zygotene stages^7,10–12^. Here, chromatin acquires a polarized-clustered, crescent shape, triggered in part by the recruitment of chromosomes ends to the nuclear periphery^13^ and in part by the destabilization of the nuclear lamina network^14^. A family of zinc finger proteins mediates the association of chromosome-specific sub-telomeric regions called Pairing Centers (PC) to the nuclear envelope, exerting an essential role for pairing and synapsis of the corresponding chromosome: ZIM-1, −2 or −3 individually bind to the PCs of the autosomes, while HIM-8 is recruited to the PC of chromosome X^15–18^. Both the PCs and their cognate PC-binding protein together with MJL-1^19^ associate with the inner nuclear envelope protein SUN-1, which becomes concentrated at the nuclear envelope associated end. This is followed by vigorous chromosome end-led movements along the nuclear periphery, essential to accomplish homologous pairing and licensing of synapsis along aligned homologous chromosomes^7,9–13^.

PCs recruit the Polo-like kinase PLK-2, which leads to SUN-1 phosphorylation and likely other targets, and is crucial for faithful pairing and synapsis^17,20^. Importantly, PLK-2 and phosphorylated SUN-1 concentrated at the PCs are part of a surveillance system that can extend the TZ stage in response to defects in synapsis and recombination^17,20,21^ and therefore these markers can be used as a proxy to monitor meiotic entry and homology search competence.

Association between coaligned homologs is stabilized by the Synaptonemal Complex (SC), a meiosis-specific proteinaceous structure composed of lateral and central elements that is conserved in most species. The SC acts as a zipper-like scaffold that allows physical exchange of DNA molecules between the homologs during recombination and thus it is essential for CO formation^22^. In worms, a family of four HORMA-domain containing proteins is recruited to the chromosome axes, where these factors play different roles during SC assembly. HTP-3 serves as a scaffold and promotes axes morphogenesis and also DSB induction^23,24^, HIM-3 is required for synapsis^25^, and HTP-1/2 play regulatory roles by coordinating homologous pairing with SC assembly^26^. Lack of HTP-3 or HIM-3 abrogates chromosome pairing, chromatin clustering and synapsis altogether, while removal of HTP-1 triggers precocious exit from the homology search state resulting in severely reduced chromosome clustering, defective pairing, and extensive non-homologous synapsis. Abrogation of *htp-2* functions on its own does not result in aberrations of pairing or synapsis, however its removal in *htp-1* mutants exacerbates CO and synapsis defects^27^, suggesting a partial redundancy.

While being an essential requirement for CO formation in *C. elegans*, the SC alone is not sufficient to promote COs, whose formation depends on homologous recombination-mediated repair of deliberately induced DNA double-strand breaks (DSBs) by the Topoisomerase-like protein Spo11^28^. Spo11 acts in conjunction with several accessory factors shown to be essential for achieving optimal DSB levels, and while Spo11 itself is highly conserved, auxiliary pro-DSB proteins have widely diverged across species^29^. Besides HTP-3 and the MRN/X complex components MRE-11 and RAD-50^30–32^, several DSB-promoting proteins have been identified in worms, including HIM-17, HIM-5, XND-1, DSB-1-2-3, but also the poly(ADP-ribose) glycohydrolase PARG-1^33–39^. Importantly, unlike in most species, DSB induction and SC formation occur independently of one another in *C. elegans*^40^, allowing the study of either process without confounding effects.

We undertook a biochemical approach to identify novel factors interacting with DSB promoting proteins, by performing mass spectrometry analysis on HIM-17 pulldowns. We found the zinc finger protein BRA-2 as a HIM-17 interactor, for which no role in meiosis has been reported yet. We recapitulated the presence of BRA-2-HIM-17 complexes *in vivo* by co-immunoprecipitation and found that the two proteins display largely overlapping localization in the germ line, yet their chromatin loading was independent. Loss of BRA-2 severely compromises fertility and reduces chiasmata formation due to a dramatic impairment of synapsis. While not being essential for SC formation or establishment of pairing on its own, removal of HIM-17 from BRA-2-depleted animals completely abolished pairing and synapsis along the autosomes; at the late pachytene stage a fully assembled SC was exclusively observed on paired X chromosomes. Strikingly, BRA-2 removal restored a transition zone nuclear morphology and recruitment of PLK-2 to the chromosome PCs in *htp-1* mutants, without improving pairing levels, lending support to BRA-2 having an early regulatory activity at the level of SC licensing. Furthermore, we discovered that removal of *him-17* in *htp-1* mutants delayed SC formation and greatly improved pairing, indicating that at least two partially redundant pathways exist in *C. elegans* to control chromosome pairing and SC formation. Our data suggest that licensing of SC assembly is regulated by BRA-2 together with HIM-17 at meiotic entry and unveil a yet undescribed role for the chromatin associated protein HIM-17 beyond DSB formation.

## RESULTS

### BRA-2, but not its paralog BRA-1, is required for robust crossover formation

In a mass spectrometry analysis aimed at identifying novel interactors of the THAP (<u>TH</u>anatos <u>A</u>ssociated <u>P</u>rotein) -domain containing protein HIM-17, we identified BRA-2 (<u>B</u>MP <u>R</u>eceptor <u>A</u>ssociated family member <u>2</u>) as a putative binding partner (Supp. Table 1), for which no roles in meiosis were known.

BRA-2 is a small protein (∼25kD) harboring a MYND-type zinc finger domain (ZnF) at its C-terminal region, conserved in several proteins across species. BLAST search highlighted the mammalian putative histone H3.3K36me3 reader ZMYND11^41^ as the closest homologue to *C. elegans* BRA-2 (Fig. 1A). In mammals, it has been shown that recognition and binding to trimethylated histone H3.3K36 by ZMNYD11 is important to regulate pre-mRNA processing and elongation, and further, mutations in human ZMYND11 have been linked to a wide spectrum of intellectual disability syndromes^42^. However, functional analyses in the germ line have never been carried out so far. It is important to note that while the MYND-ZnF domain is highly conserved between worms and mammals, the protein domains directly involved in the recognition and binding of the chromatin modification (PHD-RING, PWWP- and BROMO-domain) are not present in *C. elegans* BRA-2, possibly indicating a different function or an alternative mode of action in nematodes.

**Figure 1.**
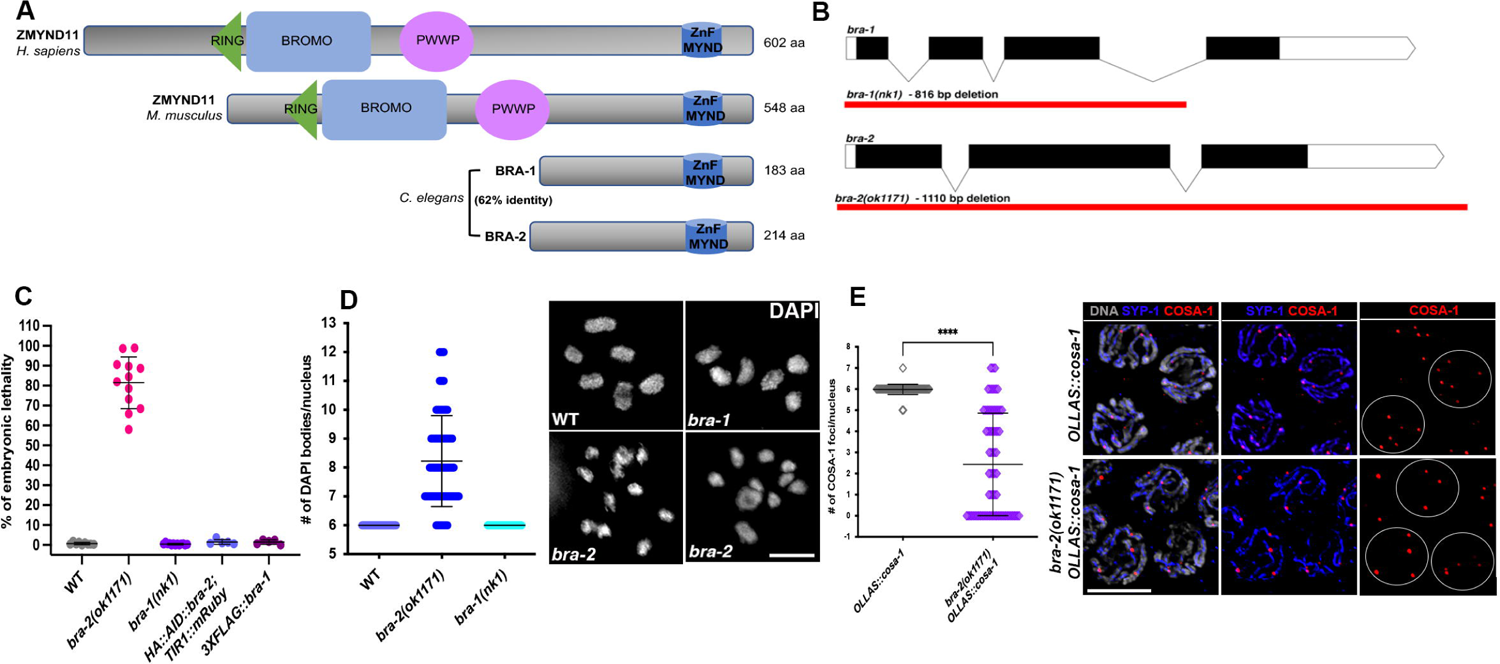
BRA-2 is evolutionarily conserved and is required for robust CO formation. **(A)** Schematic representation of *C. elegans* BRA-1/2 and mammalian ZMYND11 with putative protein domain highlighted. **(B)** Exon/Intron organization of *bra-1* and *bra-2* loci. Red lines indicate deleted regions in the respective mutant alleles. **(C)** Quantification of viability levels in the indicated genetic backgrounds. **(D)** Left: quantification of DAPI-bodies number in the indicated genotypes. Representative examples are shown to the right. Scale bar 2 μm. **(E)** Quantification of OLLAS::COSA-1 foci in the indicated genotypes (left) and representative images of late pachytene nuclei stained for SYP-1/OLLAS (COSA-1), and counterstained by DAPI. Scale bar 5 μm.

To assess the phenotypes triggered by loss of BRA-2, we exploited the *bra-2(ok1171)* mutant strain, which carries a large deletion that removes the entire *bra-2* locus, thus resulting in a null allele (Fig. 1B). Assessment of viability levels revealed high embryonic lethality in *bra-2* mutants (Fig. 1C), indicative of chromosome mis-segregation.

In nuclei at diakinesis, the last stage of meiotic prophase I, chromosomes become strongly condensed and appear as six DAPI bodies in WT animals, each representing a pair of homologous chromosomes held together by a chiasma. Defects in recombination and DNA repair can cause an increase/reduction in the DAPI bodies number and/or a variation in their morphology, thus diakinesis nuclei can be used as a powerful read out for the proper completion of upstream meiotic tasks.

Consistent with elevated embryonic lethality, diakinesis nuclei in *bra-2(ok1171)* mutants displayed a large proportion of achiasmatic univalents (89.4% of nuclei contained >6 DAPI bodies), suggesting defective CO establishment (Fig. 1D). Quantification of CO-designation sites by assessing COSA-1 foci formation^43^ revealed in fact a dramatic reduction compared to controls, with an average of 2.4 COSA-1 foci/nucleus in the *bra-2* mutants versus 5.98 in the controls (Fig. 1E), and with 40% of nuclei in the *bra-2* mutants that did not show any COSA-1 foci. This indicates that BRA-2 is essential for efficient CO formation.

The *C. elegans* genome encodes a closely related paralog of *bra-2* called *bra-1*^44^. These proteins share 62% identity and like BRA-2, also BRA-1 carries only a MYND-type ZnF domain (Fig. 1A). Given their high similarity, we sought to investigate whether BRA-1 might also be involved in promoting CO formation. We employed the *bra-1(nk1)* mutant background, which carries a large deletion that removes most of the *bra-1* locus^44^ (Fig. 1B), likely resulting in a null allele. Surprisingly, both viability levels as well as number of DAPI bodies in the diakinesis nuclei were indistinguishable from WT animals (Fig. 1C-D). This indicates that BRA-1 is not essential for chiasmata formation and that it does not act redundantly with BRA-2, suggesting that only the latter exerts essential functions during meiosis in the *C. elegans* germ line.

### BRA-2 is essential for normal SC assembly

Defects in CO formation can originate from defects in pairing, synapsis and recombination^11,26,40,45,46^. To distinguish between these possibilities, we first assayed pairing levels by performing FISH analysis using specific probes recognizing different autosomes, as well as immunostaining with antibodies directed against the PC protein HIM-8 to identify the chromosome X. We divided the gonads into five equal regions from Transition Zone (TZ) to Late Pachytene (LP) and assessed pairing of the indicated chromosomes in each nucleus across these zones (Fig. 2A). Pairing was largely achieved in *bra-2* mutants, although it peaked at slightly reduced levels compared to control animals, most prominently for the autosomes scored (Fig. 2A-B). This indicates that although not being essential, yet BRA-2 is required to attain robust homologs pairing.

**Figure 2.**
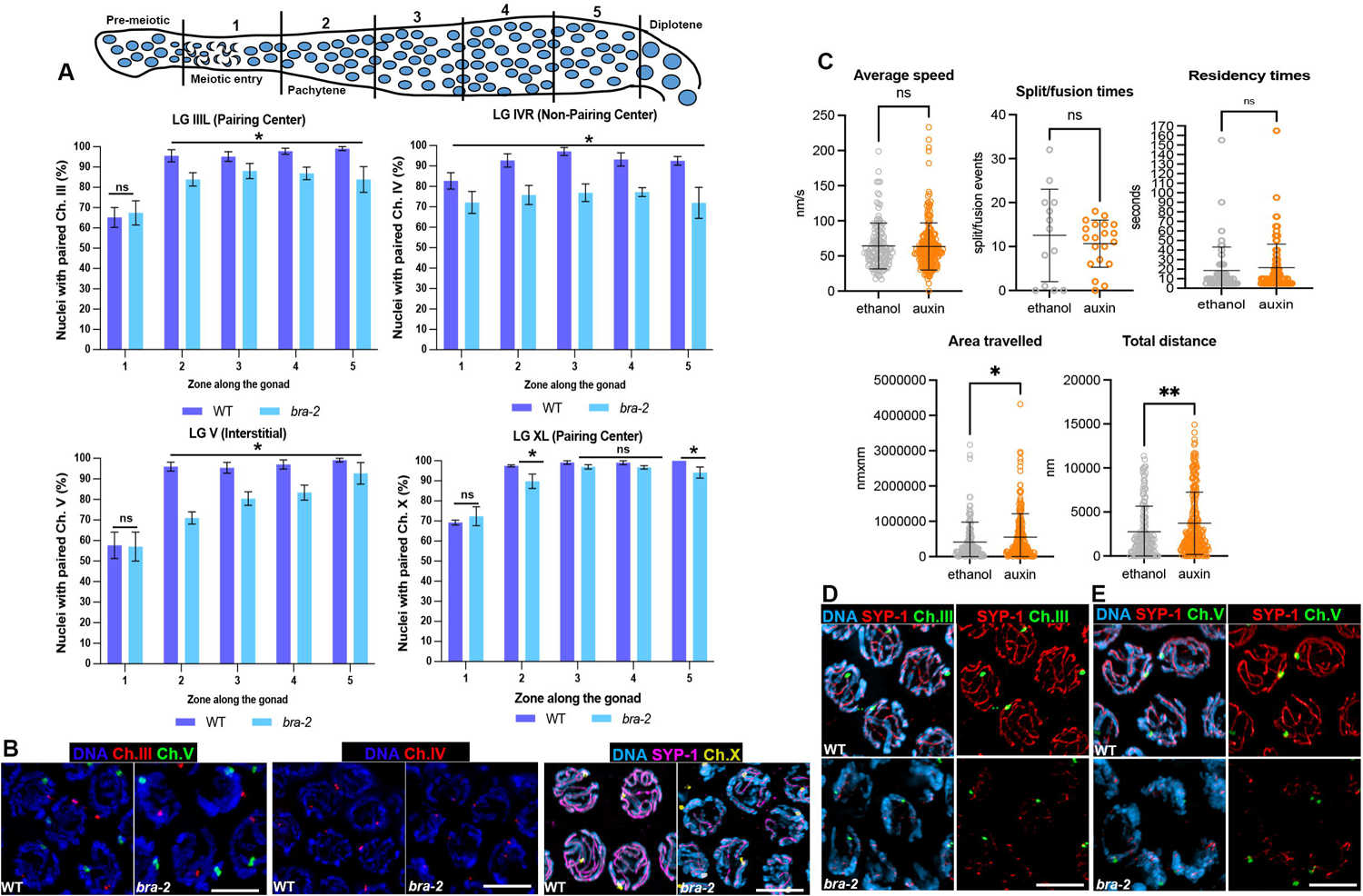
Pairing dynamics are largely unaffected in absence of BRA-2. **(A)** Schematic representation of the *C. elegans* germline with indicated Prophase I stages and zones division employed for FISH quantification. Charts show paring quantification of the indicated chromosomes in WT and *bra-2(ok1171)* mutants. Scale bars report SD and asterisk denotes statistical significance as calculated by χ*^2^* test (CI. 0.05). **(B)** Mid-pachytene nuclei displaying indicated probes signal and antibodies, counterstained by DAPI. Scale bar 5 μm. **(C)** Quantification of chromosome movement indexes in live animals assessed by tracking SUN-1::GFP in untreated *HA::AID::bra-2; TIR1::mRuby* worms and after auxin exposure. Bars indicate SD with mean. Statistical analysis was performed by two-tailed Mann-Whitney test (*ns*=non-significant, *\*p*=0.02, *\*\*p*=0.004). **(D)** Early-pachytene nuclei showing anti-SYP-1 immunostaining combined with FISH detecting chromosome III or **(E)** chromosome V. Scale bar 5 μm.

To further understand the role of BRA-2 during establishment of pairing, we investigated the movement of chromosome ends by monitoring SUN-1 dynamics in live animals, since this poses an essential requirement for successfully achieving the installation of the synaptonemal complex between homologous chromosomes^7,13^.

To this end, we engineered the *bra-2* endogenous locus by introducing an auxin-induced degradation tag (AID) in fusion with an HA tag after the start codon. This allows conditional depletion of BRA-2 upon exposure to auxin, mediated by TIR1 binding and subsequent targeting of HA::AID::BRA-2 for proteosome-dependent degradation^47^. *HA::AID::bra-2; TIR1::mRuby* animals displayed normal levels of embryonic viability (Fig. 1C), indicating that the HA::AID fusion tag did not compromise protein functionality. We assessed BRA-2 depletion efficiency by Western Blot on total protein extracts, which showed that exposure to auxin can rapidly and efficiently deplete >95% of HA::AID::BRA-2 within 3 hours (Supp. Fig. 1A). We then used auxin-depleted *HA::AID::BRA-2; TIR1::mRuby* animals carrying a functional *sun-1::GFP* transgene and tracked the dynamic behavior of GFP aggregates, which mark chromosome ends at the nuclear periphery, in live worms^12^. This analysis showed that the speed, split/fusion events and coalescence times of SUN-1::GFP aggregates followed normal kinetics in absence of BRA-2 (Fig. 2C). However, we found a mild, although statistically significant increase in the area covered and the total distance travelled by the SUN-1::GFP patches, suggesting a slightly elevated chromosome movement. These results indicate that the reduced pairing levels observed in *bra-2* mutants is not a consequence of gross defects in chromosome motion at pairing stages.

Once homology is satisfied, the physical interaction between the coaligned chromosomes is stabilized by the SC, which in *C. elegans* is assembled at meiotic entry^40,48,49^. Here, axial elements are loaded first (HTP-1/3, HIM-3, cohesins), and then central components (SYP-1/6, SKR-1/2) polymerize between the chromosome axes^23,25,48^. By early pachytene (EP), the SC is fully built along the entire length of the chromosomes, as defined by complete overlap between axial and central elements, coupled with loss of clustered chromatin configuration. DNA regions decorated by axial components but devoid of SYPs, indicate absence of synapsis.

The *bra-2(ok1171)* mutants displayed a dramatic impairment of synapsis, as SYP-1 loading between TZ and EP was barely detectable (Fig. 3A and Supp. Fig. 1B, mostly in puncta and small patches). In MP-LP stages (corresponding to zones 5-6 in the gonad), SYP-1 staining appeared more robust and elongated, however full synapsis was only rarely observed, indicating that BRA-2 is essential for normal SC assembly. Furthermore, the chromatin in TZ nuclei displayed a clustered morphology and appropriate HTP-3 localization in *bra-2* mutants, suggesting successful meiotic entry and a genuine synapsis defect. We also analyzed synapsis in *bra-1* and *bra-2; bra-1* double mutants by HTP-3/SYP-1 immunostaining, which showed no dramatic defects in the former and no differences to *bra-2* single mutants in the latter (Fig. 3A and Supp. Fig. 1B), further corroborating that only BRA-2 exerts meiotic functions. Lastly, by combining FISH analysis (or HIM-8 detection) with SYP-1 immunostaining, we were able to observe that paired chromosomes X, III and V and were always associated with some SYP-1 protein (Fig. 2B, 2D-E), denoting that the little residual installation of SC does not occur between non-homologous chromosomes in absence of BRA-2.

**Figure 3.**
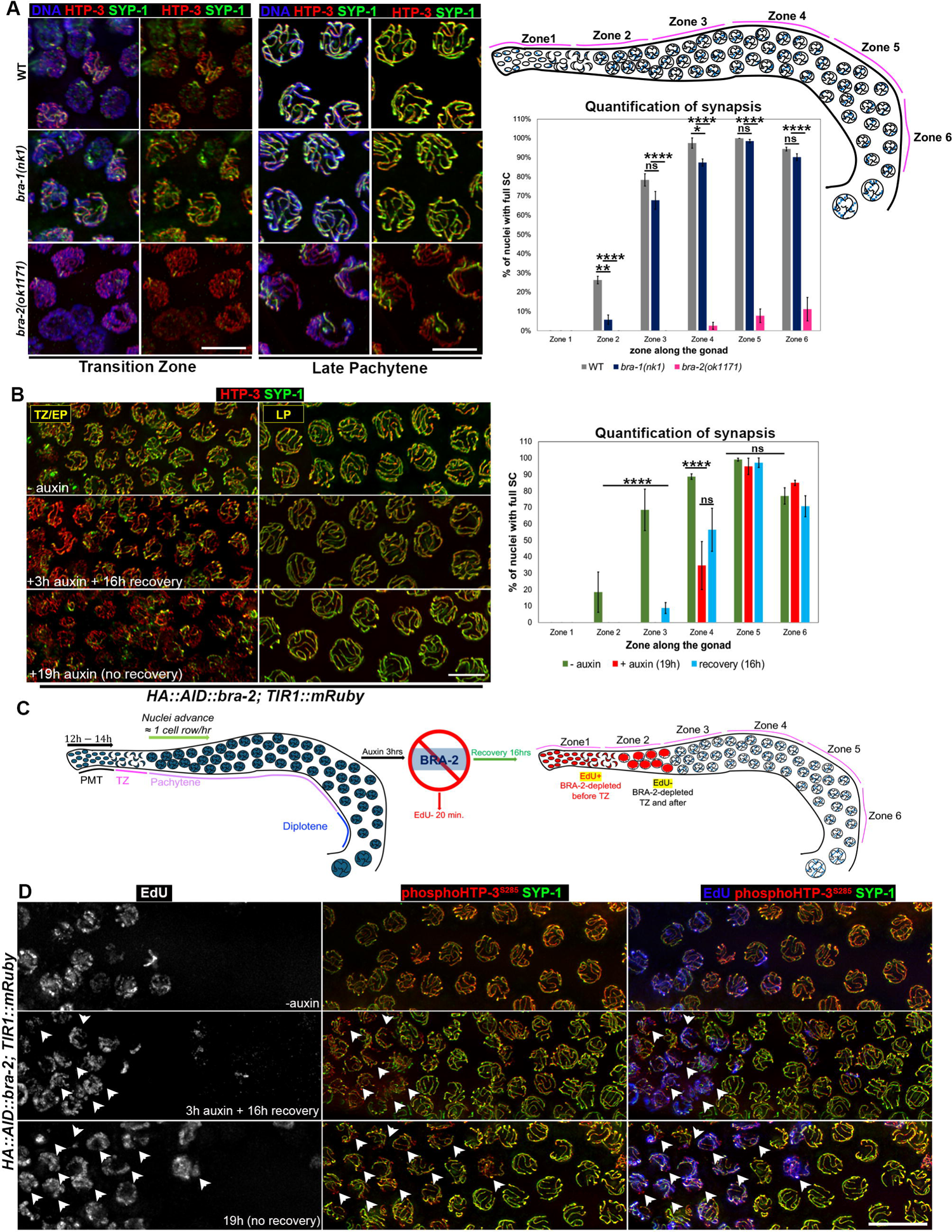
BRA-2 is essential for normal establishment of synapsis. (A) **<u>Left:</u>** nuclei from the indicated mutants and stage stained for axial (HTP-3) and central (SYP-1) elements of the SC. Note extensive regions of HTP-3 devoid of SYP-1 signal, indicating absence of synapsis. Scale bar 5 μm. <u>Right:</u> schematic representation depicting zoning of the *C. elegans* germ line used to quantify synapsis. Chart shows quantification of nuclei with full SC in the indicated zones and genetic backgrounds. Bars show S.E.M. and asterisks denote statistical significance assessed by two-tailed Mann-Whitney test (*ns=non-significant*, *\*\*p=0.004*, *\*\*\*\*p<0.0001*). **(B)** Left: representative images of nuclei from *HA::AID::bra-2; TIR1::mRuby* animals at the indicated stages and conditions of exposure to auxin. Scale bar 5 μm. Right: Quantification of nuclei displaying full synapsis across the germ line in the indicated backgrounds and exposure conditions to auxin. Bars show S.E.M. and asterisks denote statistical significance assessed by two-tailed Mann-Whitney test (*ns=non-significant*, *\*\*\*\*p<0.0001*). **(C)** Schematization of the assay performed to track stage-specific cells before and after BRA-2 depletion. **(D)** Co-staining of SYP-1/pHTP-3^S285^ with EdU in the indicated conditions of exposure to auxin. Arrowheads point to nuclei that are positive for EdU incorporation and show incomplete SC. Scale bar 10 μm.

### Loss of BRA-2 destabilizes SC subunits and hinders their polymerization

We next considered the possibility that impaired SC assembly observed in *bra-2* mutants could be resulting from defective cell-cycle progression.

To address this, we co-stained WAPL-1 and the Cyclin E CYE-1, two markers expressed in the progenitor zone (which harbors nuclei in premeiotic divisions and pre-meiotic replication)^50,51^ (Supp. Fig. 2A). Moreover, we performed an assay to monitor incorporation of 5-Ethynyl-2′-deoxyuridine (EdU) to assess ongoing replication (Supp. Fig. 2B). For all these markers we did not find any difference between WT and *bra-2* mutants, indicating that germ cells enter meiosis with unperturbed kinetics and undergo normal mitotic and pre-meiotic replication respectively.

Once we assessed that the mitosis-meiosis developmental switch was not impacted by lack of BRA-2, we combined our *HA::AID::bra-2; TIR1::mRuby* strain with available functional tagged lines of other SC central elements, to monitor loading of SYP-2::V5, V5::SYP-4^52^, and GFP::SYP-3^9^.

*HA::AID::bra-2; TIR1::mRuby* animals exposed to auxin for 24 hours recapitulated the synapsis defects observed in the *bra-2(ok1171)* null mutants, whereby almost no SYP-1 loading was observed until EP stage (Supp. Fig. 2C). We found that similar to defective SYP-1 loading, BRA-2-depleted animals also displayed impaired localization of SYP-2::V5 (Supp. Fig. 2D), GFP::SYP-3 (Supp. Fig. 2E) and V5::SYP-4 (Supp. Fig. 2F), as expected by their interdependent localization. Interestingly, Western blot analysis showed that SYP-2::V5 and V5::SYP-4 were destabilized in absence of BRA-2 (Supp. Fig. 2G) despite normal transcription levels (Supp. Fig. 2H), which prompted us to investigate whether the initial loading or rather the polymerization of the SC central elements along the axes was compromised.

To distinguish between these two possibilities, we carried out live imaging of GFP::SYP-3 at meiosis onset in live worms^9^, which revealed that the initial recruitment of SYP-3 as nucleation sites (bright foci) was slightly reduced (Supp. Fig. 2I). However, while in presence of BRA-2 (-auxin) GFP::SYP-3 foci underwent elongation, we found that only about half of detected nucleation events showed elongation upon BRA-2 depletion (Supp. Fig. 2J). However, these displayed a severe reduction in the elongation rate of about 65% (474nm/m in the -auxin vs 169nm/m in the +auxin) (Supp. Fig. 2K-L). Altogether, these results indicate that BRA-2 promotes the efficient recruitment of the SC central elements upon meiotic entry as well as their optimal polymerization along the axes.

### BRA-2 promotes licensing rather than maintenance of synapsis

We observed that following exposure to auxin for 24 hours nuclei at meiotic entry displayed similar SC defects to those observed in *bra-2* null animals, however, those at later stages (LP) remained fully synapsed (Supp. Fig. 2M).

Germ cells advance synchronously from the distal tip of the gonad onwards, with a pace of approximately one nuclear row/hour ^53,54^. Therefore, we reasoned that impaired synapsis in nuclei specifically at TZ-EP would be consistent with BRA-2 exerting more prominent roles at earlier rather than later stages. In support of this, we observed that if exposure to auxin was prolonged for 48 hours, thus allowing meiocytes to travel further within the gonad, then synapsis defects were in fact detected throughout pachytene, fully phenocopying the *bra-2* null mutants (Supp. Fig. 3A).

To further corroborate this, we undertook two different approaches: first, we exposed *HA::AID::bra-2; TIR1::mRuby* animals to auxin for 3 hours to achieve nearly complete BRA-2 depletion in the whole germ line (Supp. Fig. 1A) and then we moved the animals onto plates without auxin to allow for BRA-2 resynthesis. We reasoned that analyzing the depleted animals at different times during recovery/resynthesis would allow us to assess how cells that were at different substages of meiotic progression during auxin exposure responded to BRA-2 depletion.

Given that it takes 12-14 hours for the nuclei to travel from the mitotic tip to TZ^53–56^, we let the depleted animals recover for slightly longer (16 hours) to ensure that cells had enough time to enter meiosis. As shown in Fig. 3B, nuclei in zone 3 and 4 (corresponding to TZ/EP), that were in the premeiotic tip/meiotic entry at the time of the depletion, displayed comparable SC impairment to worms constantly exposed to auxin without recovery. Nuclei in zone 5 and 6 (corresponding to MP/LP), depleted for BRA-2 when they were already beyond TZ, displayed instead comparable levels of synapsis as control animals.

In a second approach, we treated the *HA::AID::bra-2; TIR1::mRuby* animals as above but we included incubation with EdU during the last 20 minutes of the 3h-exposure to auxin, followed by the 16h of recovery. Since the EdU is incorporated into the DNA only during premeiotic DNA replication, this allowed us to clearly distinguish the germ cells that engaged meiosis after depletion of BRA-2 (EdU-positive) from those that had entered meiosis before/during the depletion (EdU-negative) (Fig. 3C). As shown in Fig. 3D, the cells displaying unsynapsed chromosomes were also positive for EdU (arrowheads) and conversely, EdU-negative cells showed full synapsis. Importantly, Western blot analysis showed that significant resynthesis of BRA-2 protein levels occurred within 24-48h (Supp. Fig. 3B), ruling out any major contribution as a result of its reappearance and resumption of its function within the 16 hours of the recovery time window.

Altogether, these data indicate that BRA-2-mediated regulation of chromosome synapsis occurs at meiotic entry and lends support to our initial hypothesis that BRA-2 is essential for the timing of SC initiation rather than its maintenance.

### Loss of BRA-2 restores chromosome clustering and loading of PLK-2 at the nuclear periphery in htp-1 mutants

Homologous pairing and SC assembly occur as early meiotic events, and previous work has shown a crucial involvement for the HORMA-domain containing protein HTP-1 in coordinating these processes^26,57^. Furthermore, analyses in pairing-defective, but synapsis-proficient mutants, have also shown that the SC can be polymerized irrespectively of homology in *C. elegans*^11,58,59^, indicating that tight controlling mechanisms exist to ensure proper coordination of pairing with the establishment of homologous synapsis. Given the early requirements imposed by *bra-2* functions on SC assembly, we wondered about its epistatic relationship to *htp-1*.

To this end, we combined the *HA::AID::bra-2; TIR1::mRuby* animals with *htp-1(gk174)* null mutants and assessed PLK-2/SYP-1 stainings.

PLK-2 undergoes a dynamic localization in the germ line, as it concentrates into aggregates at the nuclear periphery during TZ stage, and it redistributes along the SC in pachytene in response to CO site designation and to promote timely inactivation of CHK-2 ^17,20,60^. Importantly, PLK-2, together with phosphorylated SUN-1, can delay exit from TZ (visible as prolonged nuclear clustering) in the presence of defects in synapsis or recombination^17,20,21,57^.

Abrogation of *htp-1* function triggers precocious exit from TZ and extensive non-homologous synapsis, therefore *htp-1* null mutants lack a defined region of the gonad containing nuclei with clustered chromatin^26^ and display a dramatic reduction of PLK-2/pSUN-1^S^^8^ at the nuclear envelope^57^.

PLK-2 aggregates at the nuclear periphery were barely detectable at meiotic entry in *htp-1* mutants (TZ, -auxin, Fig. 4A, and Supp. Fig. 4), and no clearly defined nuclear polarization was observed, recapitulating previous findings^57^. Exposure of *HA::AID::bra-2; TIR1::mRuby* animals to auxin triggered retention of PLK-2 at the PCs (TZ, +auxin, Fig. 4A), in line with lack of synapsis. Strikingly, removal of BRA-2 in *htp-1* mutants restored chromatin clustering, which consistently overlapped with persistent PLK-2 localization at the nuclear envelope until MP stage (TZ and MP, +auxin, Fig. 4A and Supp. Fig.4). SYP-1 loading appeared greatly diminished compared to both BRA-2-depleted worms and *htp-1* single mutants, as only one or two SYP-1 tracks were detectable in MP nuclei of worms lacking both BRA-2 and HTP-1 (Fig. 4B). This indicates that lack of BRA-2 largely abrogates indiscriminate SC assembly triggered by absence of HTP-1 and restores establishment and maintenance of chromosome clustering.

**Figure 4.**
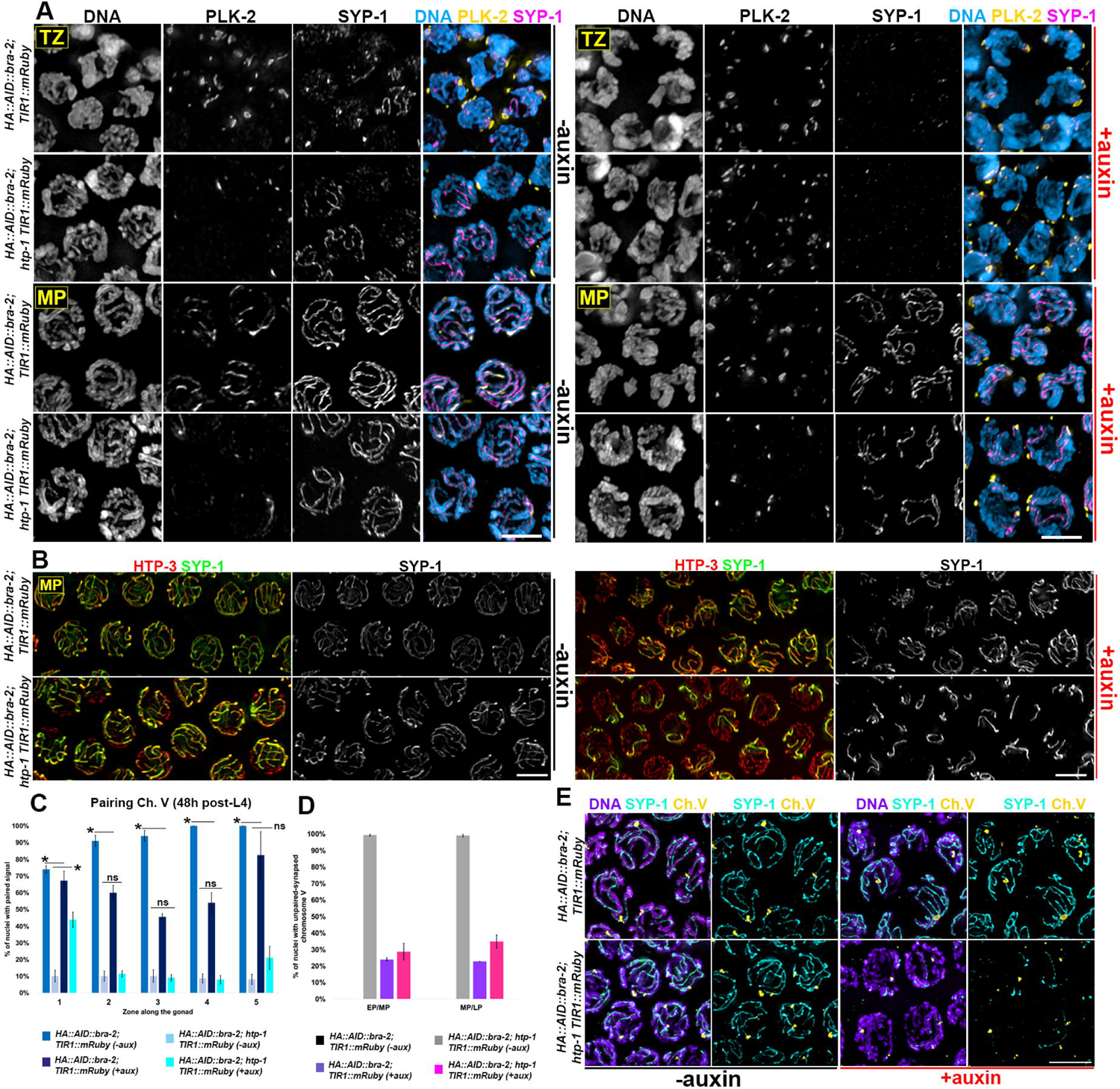
Loss of BRA-2 restores nuclear clustering and PLK-2 loading in *htp-1* mutants. **(A)** Nuclei in transition zone (TZ) or mid-pachytene (MP) stained for PLK-2 and SYP-1 in *HA::AID::bra-2; TIR1::mRuby* control animals and *htp-1(gk174)* mutants, before and after auxin exposure (24h). Scale bar 5 μm. **(B)** Mid-late pachytene nuclei stained for HTP-3/SYP-1 in the indicated genetic backgrounds before and after exposure to auxin. Scale bar 5 μm. **(C)** Quantification of nuclei with paired signals for Ch. V in the indicated genetic backgrounds before and after exposure to auxin. Gonads were divided into five equal regions spanning TZ-LP. Bars depict S.E.M. and asterisks indicate statistical significance as assessed by χ^2^ test (CI=95%). **(D)** Quantification of non-homologous synapsis in the indicated genetic backgrounds and exposure conditions to auxin only in the pachytene region. Bars indicate S.E.M. **(E)** Representative images of mid-pachytene nuclei showing FISH for Ch. V combined with anti-SYP-1 immunostaining. Scale bar 5 μm.

Given that chromatin clustering results from vigorous chromosome movement, which in turn is essential for robust synapsis between the homologous chromosomes, we wondered whether the rescue of TZ coupled with efficient PLK-2 recruitment at the nuclear envelope were also indicative of improved pairing levels in the *htp-1* mutants depleted for BRA-2.

To this end, we performed FISH analysis to monitor pairing of chromosome V. To make sure that also nuclei in LP were affected by BRA-2 depletion, we exposed the *HA::AID::bra-2; htp-1 TIR1::mRuby* animals to auxin for 48h instead of 24h, as we have previously shown that at this time point also cells at later stages display SC defects (Supp. Fig. 3A).

While initial pairing of chromosome V was slightly improved (zone 1 - 43.8% in BRA-2-depleted *htp-1* mutants versus 10.1% in non-depleted), removal of BRA-2 did not bypass the constraint imposed by *htp-1* function in achieving robust chromosome coalignment in the rest of the germ line (Fig. 4C), since the percentage of nuclei with unpaired chromosomes from zone 2-6 was comparable to *htp-1* mutants. We also noticed that in the *HA::AID::bra-2; TIR1::mRuby* animals, there was a higher frequency of nuclei with unpaired signals compared to *bra-2(ok1171)* nulls (Fig. 2). We hypothesize that this might be a consequence of the age difference of the worms analyzed between the two experiments, as worms were analyzed 24h post-L4 when the *bra-2(ok1171)* deletion mutant was used, versus 48h post-L4 in the *bra-2* degron strain. When we combined FISH with SYP-1 immunostaining we found that the chromosome V was engaged in non-homologous synapsis (assessed by monitoring whether one or both probe signals were associated with SYP-1) in all pachytene nuclei scored in the *htp-1* null mutants, recapitulating previous data^26,27^. In contrast, these were reduced to levels observed in control animals upon BRA-2 depletion (Fig. 4D-E), consistent with the fact that BRA-2 removal abrogates indiscriminate synapsis triggered by *htp-1* loss of function.

Altogether, these results indicate that removal of BRA-2 suppresses untimely SC polymerization caused by loss of *htp-1* function but is not sufficient to bypass HTP-1-dependent pairing.

### BRA-2 is enriched on the autosomes and its loading is independent of synapsis and recombination

To gain further insight into BRA-2 functions, we exploited our *HA::AID::bra-2* tagged line to analyze its localization in the germ line. Immunostaining using anti-HA antibodies showed that BRA-2 localizes as a nuclear factor throughout meiotic prophase I (Fig. 5A). This nuclear localization was recapitulated by Western blot on fractionated protein extracts, which indeed showed that HA::AID::BRA-2 is enriched in both soluble and chromatin-bound nuclear fractions (Fig. 5B). Cytological analysis showed that HA::AID::BRA-2 was not evenly distributed on chromatin, as one chromosome was devoid of signal. Co-staining using anti-HIM-8 antibodies revealed that HA::AID::BRA-2 is prominently excluded from the chromosome X (Fig. 5C), as also indicated by colocalization with H3K4me2, XND-1 and phosphorylated-RNA Pol II^S^^2^ (Supp. Fig. 5A), all known to be enriched on the autosomes^35,61,62^. Furthermore, assessment of HA::AID::BRA-2 localization in the *spo-11(ok79)* and *syp-2(ok307)* mutants showed that loading of BRA-2 is independent of DSBs and synapsis respectively (Supp. Fig. 5B).

**Figure 5.**
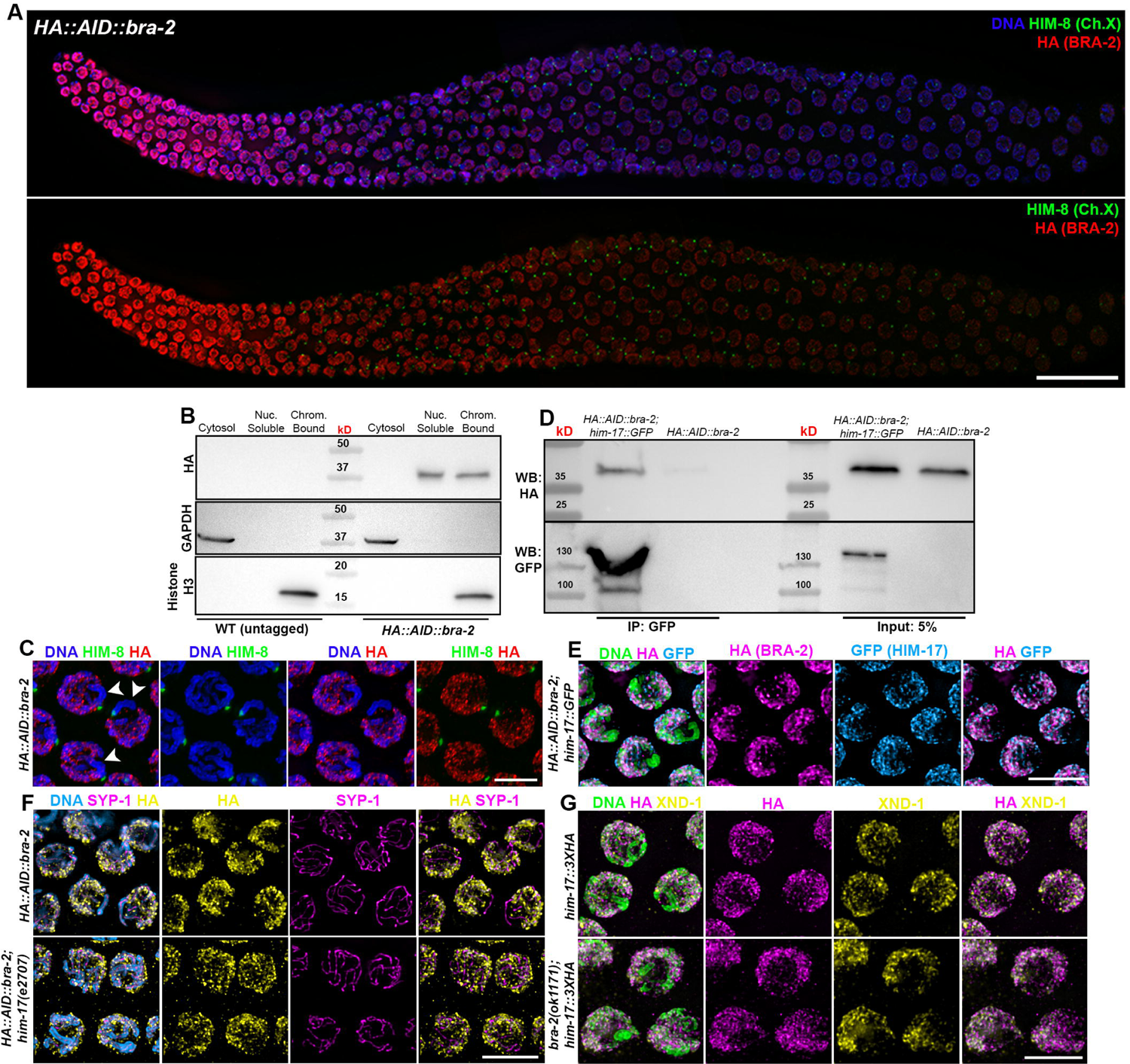
BRA-2 is enriched on the autosomes and physically interacts with HIM-17. **(A)** Top: whole mount gonad stained with anti-HA and anti-HIM-8, showing expression of BRA-2 throughout the germ line and its enrichment on the autosomes. Scale bar 20 μm. **(B)** Western blot on fractionated protein extracts probed with anti-HA antibodies showing enrichment of BRA-2 in the nuclear-soluble and chromatin-bound fractions. Detection of GAPDH and Histone H3 was used as loading controls of cytosolic and chromatin-bound fraction respectively. **(C)** Magnified mid-pachytene nuclei showing more detailed staining of HA::AID::BRA-2/HIM-8. Arrowheads indicate the X chromosome. Scale bar 5 μm. **(D)** Co-immunoprecipitation assay showing physical interaction between HIM-17::GFP and HA::AID::BRA-2. **(E)** Early pachytene nuclei stained for GFP (HIM-17) and HA (BRA-2) showing extensive overlapping localization. Scale bar 5 μm. **(F)** Early-mid pachytene nuclei immunoassayed for HA and SYP-1 displaying *him-17*-independent recruitment of BRA-2 onto the DNA. Scale bar 5 μm. **(G)** Early-mid pachytene nuclei immunoassayed for HA and XND-1 displaying *bra-2*-independent recruitment of HIM-17 onto the DNA. Scale bar 5 μm.

Although we did not observe chromosome segregation defects in the *bra-1(nk1)* mutants nor epistatic relationship between *bra-1* and *bra-2* (Fig. 1C-D and Supp. Fig. 1), we wished to investigate on BRA-1 loading nonetheless and monitor eventual changes in absence of BRA-2. To this end, we generated a *3XFLAG::bra-1* tagged line (Fig. 1C) and combined it with *HA::AID::bra-2* to analyze their localization.

HA::AID::BRA-2 and 3XFLAG::BRA-1 display a distinct temporal loading pattern in the germ line, as the former is expressed throughout the gonad and the latter appears only at pachytene exit (Supp. Fig. 5C). BRA-1 and BRA-2 did not display interdependent loading, as 3XFLAG::BRA-1 was normally loaded in BRA-2-depleted animals and conversely, HA::AID::BRA-2 did not display any abnormalities in the *bra-1(nk1)* knockout mutants (Supp. Fig. 5C). Lastly, Western blot analysis on whole cell extracts showed that BRA-1/2 proteins are not destabilized or reduced in the reciprocal mutant backgrounds (Supp. Fig. 5D-E), further corroborating that these two factors act in distinct pathways and that BRA-1 is likely to exert negligible or no roles at all in the germ line.

Given the putative interaction identified by mass spectrometry (Supp. Table 1), we next sought to investigate whether BRA-2 and HIM-17 interact *in vivo,* and if their loading is affected in the reciprocal mutant background. Immunoprecipitation of HIM-17::GFP^33^ pulled down HA::AID::BRA-2 (Fig. 5D), confirming our mass spectrometry data and indicating that they are found in a complex. Cytological analysis showed that BRA-2 and HIM-17 display a very similar localization and largely colocalize in the gonad (Fig. 5E). However, we observed that BRA-2 was still loaded in *him-17(e2707)* mutants (Fig. 5F) and conversely, HIM-17::3XHA appropriately localized in the *bra-2(ok1171)* mutants (Fig. 5G). Thus, while displaying a largely overlapping localization and being found together in protein complexes, BRA-2-HIM-17 do not undergo interdependent loading.

### BRA-2 and HIM-17 promote synapsis of autosomes in a partially redundant manner

To address whether *bra-2* and *him-17* would cooperate, we combined the *HA::AID::bra-2; TIR1::mRuby* with the *him-17(e2707)* mutant to analyze SC assembly.

Previous studies have shown that while being important for DSB induction, HIM-17 is dispensable for pairing and synapsis^33^ and we similarly did not find detectable aberrations in SC assembly in the single mutant (Fig. 6A).

**Figure 6.**
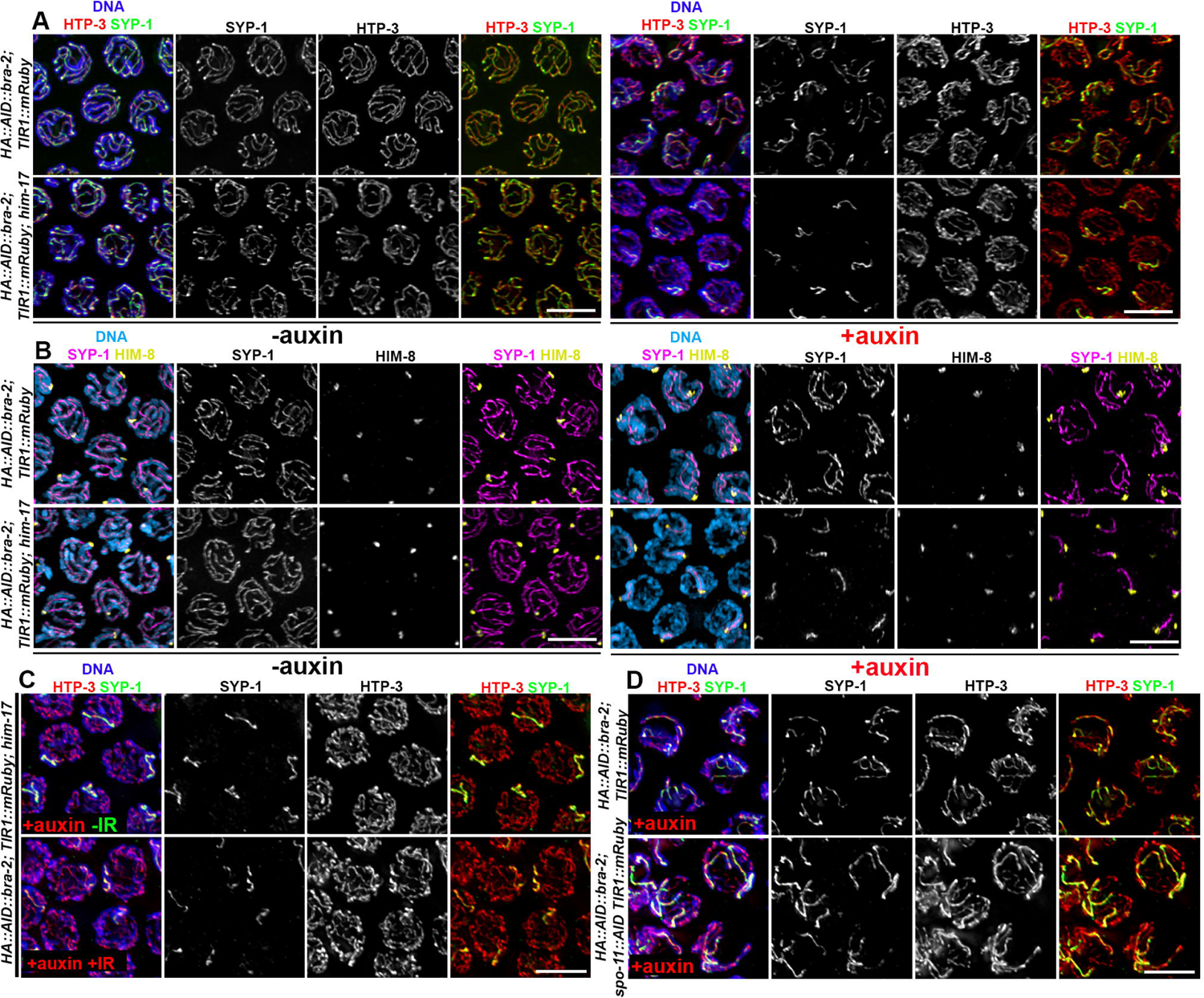
Loss of BRA-2 reveals cryptic roles for HIM-17 in promoting autosome synapsis. **(A)** Analysis of synapsis in early-mid pachytene nuclei of *HA::AID::bra-2; TIR1::mRuby* control animals and in *him-17(e2707)* mutants, before and after exposure to auxin. Scale bar 5 μm. **(B)** Co-staining of SYP-1-HIM-8 reveals that only the X chromosome is proficient in SC assembly upon contemporary absence of BRA-2 and HIM-17. Scale bar 5 μm. **(C)** Mid-pachytene nuclei of worms of the indicated genotype exposed to auxin before and after IR showing that lack of synapsis does not depend on DSBs. Scale bar 5 μm. **(D)** Mid-pachytene nuclei of worms of the indicated genotype exposed to auxin showing that lack of synapsis does not depend on SPO-11 activity. Scale bar 5 μm.

Strikingly, concomitant depletion of BRA-2 and HIM-17 entirely abrogated synapsis between TZ and mid-pachytene, and even at later stages, SYP-1 was detected mostly as a single stretch, possibly suggesting synapsis of one chromosome pair only (Fig. 6A and Supp. Fig. 6A). Co-staining using anti-HIM-8 antibodies indeed revealed that the chromosome X was the only one retaining synapsis in *him-17* mutants depleted for BRA-2 (Fig. 6B, Supp. Fig. 6B), indicating that HIM-17 promotes residual synapsis along the autosomes in absence of BRA-2, and that they are both dispensable for SC establishment along the sex chromosomes.

Importantly, we ruled out that lack of synapsis originated from impaired meiotic entry by co-staining WAPL-1 and phosphorylated SUN-1^S8^, as the former is lost from the nucleus and the latter appears at the nuclear envelope upon entry into meiosis respectively. WAPL-1 and pSUN-1^S8^ displayed mutually exclusive localization before and after exposure to auxin (Supp. Fig. 6C), further indicating that impaired SYP-1 loading is not caused by a defective mitosis-meiosis switch.

It has been previously shown that *him-17* has epistatic relationships with two other pro-DSB factors in *C. elegans*, namely *xnd-1* and *him-5*^34^. Thus, we wondered whether the synthetic synapsis defects observed under simultaneous removal of BRA-2-HIM-17 could be similarly triggered by lack of BRA-2 and these proteins as well.

Removal of HIM-5 or XND-1 in BRA-2-depleted animals did not further impair synapsis (Supp. Fig. 6D-E), indicating a specific role for *him-17* in promoting SC assembly in absence of BRA-2.

We next sought to investigate whether the impaired synapsis observed under simultaneous removal of HIM-17 and BRA-2 was linked to reduced DSB formation, since *him-17* activity has been shown to be essential for break induction^33^.

First, we exposed worms lacking both HIM-17 and BRA-2 to ionizing radiations to assess whether exogenous DSBs could suppress SC defects; and performed analysis of HTP-3 and SYP-1 staining 4 hours after irradiation, which did not reveal any differences compared to non-irradiated BRA-2-depleted animals and indicating that SC defects are independent of DSB formation (Fig. 6C).

As a complementary approach, we simultaneously depleted SPO-11 and BRA-2, as we reasoned that if lack of meiotic DSBs caused synapsis defects in the *bra-2; him-17* then this phenotype should be recapitulated in the *bra-2; spo-11*. In this case, synapsis defects were also indistinguishable from animals depleted only for BRA-2 (Fig. 6D), indicating that the synapsis defects occurred independently of DSBs *per se*.

These results indicate that HIM-17 exerts roles in promoting SC assembly which are not linked to its functions in mediating DSB induction, unveiling a previously unknown role for this protein in promoting synapsis.

### Loss of HIM-17 partially abrogates pairing and synapsis defects in htp-1 mutants

Unlike *bra-2*, *him-17* is dispensable for synapsis. However, the synthetic effect that we observed when BRA-2 and HIM-17 were simultaneously removed, might suggest that HIM-17 could be part of an accessory/parallel pathway that is activated only when the establishment or regulation of synapsis is already compromised.

To investigate this possibility, we assessed whether removal of *him-17* in *htp-1* mutants altered SC dynamics and/or pairing as similarly observed upon BRA-2 depletion (Fig. 4).

Indeed, *htp-1; him-17* double mutants showed a well-defined TZ with nuclei displaying a clustered chromatin morphology (Fig. 7A). This was coupled with reduced SYP-1 loading, which localized predominantly as chromosome-associated puncta at meiotic entry/EP, and wild-type like clusters of PLK-2-labelled chromosome ends at the nuclear periphery (Fig. 7B). Compared to *htp-1* animals depleted for BRA-2 though, the window of PLK-2 localization at the nuclear envelope was narrower in the *htp-1; him-17* doubles, extending to a size comparable to WT animals (Fig. 7C).

**Figure 7.**
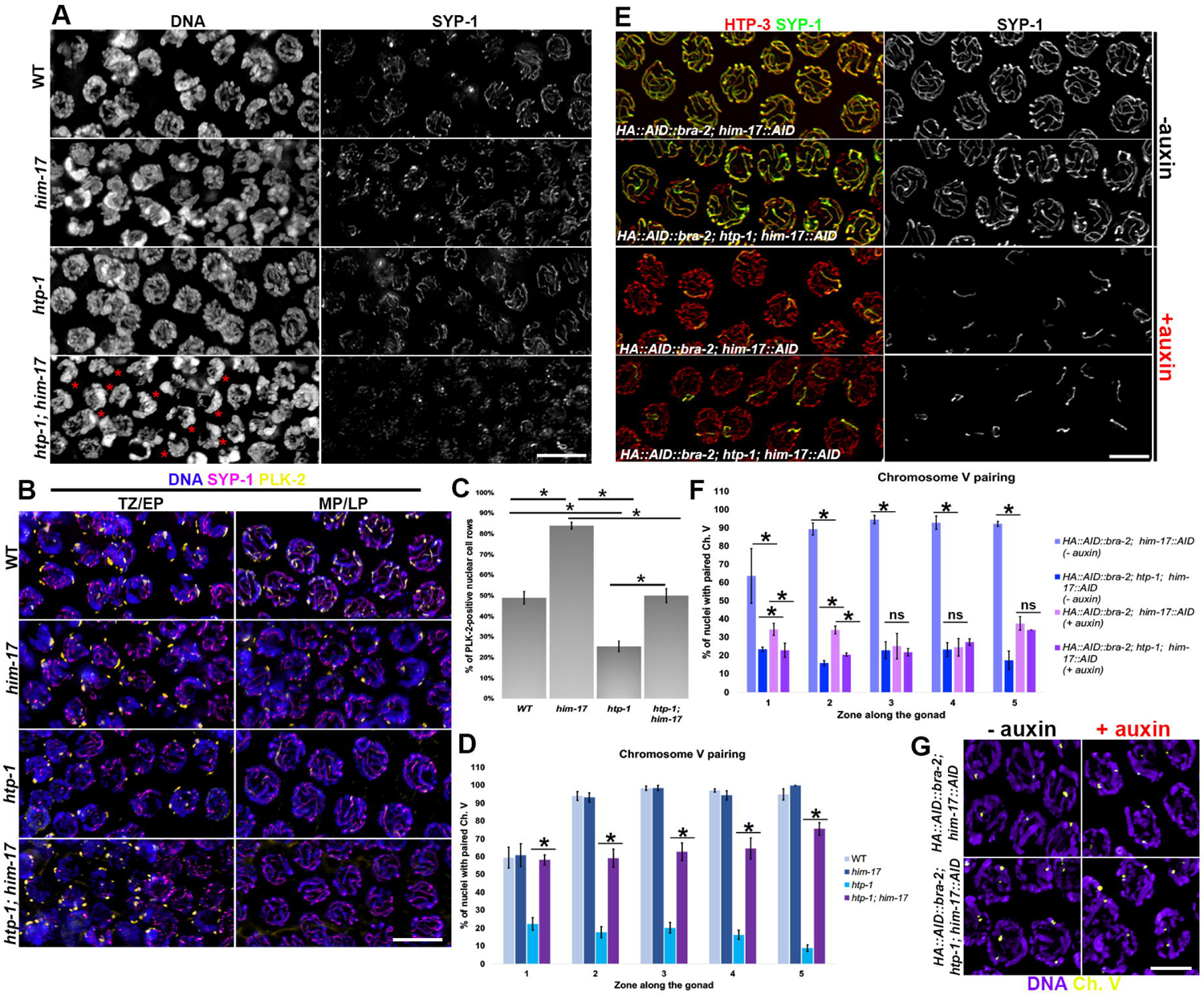
Loss of HIM-17 improves pairing in *htp-1* null mutants but abrogates homolog co-alignment in BRA-2-depleted animals. **(A)** Analysis of SYP-1 localization in the indicated mutant backgrounds. Red asterisks indicate nuclei with clustered chromatin. Scale bar 5 μm. **(B)** SYP-1/PLK-2 immunofluorescence in the indicated mutants and stages. (TZ= Transition Zone, EP=Early Pachytene, MP=Mid-Pachytene and LP=Late-Pachytene). Scale bar 10 μm. **(C)** Quantification of PLK-2 in the genotypes shown in (B). Bars indicate S.E.M. and asterisks show statistical significance assessed by T test. **(D)** Quantification of Ch. V pairing in the indicated mutants. Gonads were divided into five equal regions spanning TZ-LP. Bars indicate S.E.M. and asterisks denote statistical significance as assessed by χ^2^ test. **(E)** Mid-pachytene nuclei stained for HTP-3/SYP-1 in the indicated genetic backgrounds and conditions of exposure to auxin. All strains analyzed carried the *TIR1::mRuby* expressing transgene in the background. Scale bar 5 μm. **(F)** Quantification of Ch. V pairing in the indicated genetic backgrounds and exposure conditions to auxin. All strains analyzed carried the *TIR1::mRuby* expressing transgene in the background. Gonads were divided into five equal regions spanning TZ-LP. Bars indicate S.E.M. and asterisks denote statistical significance as assessed by χ^2^ test. **(G)** Representative images of mid-pachytene nuclei from the indicated genetic backgrounds and exposure conditions to auxin displaying FISH signal for Ch. V (quantified in F). Scale bar 5 μm.

Strikingly, FISH analysis revealed that in the *htp-1; him-17* double mutants the pairing of chromosome V in TZ (corresponding to Zone 1) was not different from the controls, and it was steadily maintained throughout the germ line (Fig. 7D, Supp. Fig. 7A). Combined analysis of FISH and immunostaining for SYP-1 also showed that paired chromosome V was normally synapsed (Supp. Fig. 7B). Thus, our data suggest that removal of HIM-17 in *htp-1* mutants not only restores chromosome clustering early on, but also allows the homologs to align and remain synapsed thereby preventing non-homologous synapsis.

The pairing and synapsis phenotypes observed in the *htp-1; him-17* worms differed from those observed in the *bra-2; htp-1* animals, since in the former we found only a delayed SYP-1 loading associated with strongly improved pairing levels, while in the latter, synapsis was almost entirely abrogated but pairing levels were not different from *htp-1* single mutants (Fig.4). Therefore, we decided to deplete both BRA-2 and HIM-17 to assess their epistatic relationships in influencing pairing and synapsis establishment in germ lines lacking *htp-1*.

To this end, we generated a degron-tagged line of *him-17* that was functional, as *him-17::AID; TIR1::mRuby* animals displayed cytologically detectable HIM-17 and abundant RAD-51 foci formation, indicating normal expression and loading of HIM-17::AID and proficiency for DSB induction (Supp. Fig. 7C).

Importantly, the BRA-2-HIM-17 double depletion recapitulated the synapsis impairment observed in the *him-17* mutants depleted for BRA-2 only (Fig.7E, Fig. 6 and Supp. Fig. 6), indicating that the double depletion results in the predicted phenotype. SC impairment was unchanged by removal of *htp-1* (Fig. 7E), suggesting that BRA-2-HIM-17-mediated roles in regulating synapsis occur irrespectively of HTP-1 presence.

Unexpectedly, FISH analysis for chromosome V revealed a dramatic impairment of pairing in the BRA-2-HIM-17 depleted animals, which was comparable to *htp-1* single mutants (Fig. 7F-G). We performed cytological analysis using antibodies directed against phosphorylated SUN-1^S8^ which showed abundant phosphorylation at the nuclear envelope and extended TZ in *HA::AID::bra-2; TIR1::mRuby, him-17(ok424)* mutants (Supp. Fig. 7D), indicating that the pairing defects did not originate from impairment of the chromosome movement machinery and that the signaling maintaining active chromosome clustering is intact, irrespectively of HTP-1 presence. These results suggest that the lack of synapsis observed in the absence of *bra-2* and *him-17* is most likely due to the inability of the homologs to successfully pair rather than to a synthetic effect that impedes SC initiation.

## DISCUSSION

Our work identifies the MYND-ZnF domain containing protein BRA-2 as a novel HIM-17 interactor and shows that it exerts crucial functions in regulating SC licensing. Lack of BRA-2 impairs synapsis without affecting homolog pairing, resulting in defective CO establishment and reduced chiasmata formation. Loss of BRA-2 reduces the polymerization rate of SC central components concomitant with protein destabilization (despite normal transcription rates). Conditional degron-mediated depletion of BRA-2 reveals that the key function of the protein during synapsis is in regulating SC licensing rather than SC stabilization, and strikingly, co-depletion with HIM-17 reveals a novel, intersecting network coordinating homolog pairing with establishment of synapsis.

### BRA-2 as a relay for SC licensing upon homology assessment

At meiotic entry, homologous chromosomes must pairwise align to successfully undergo synapsis and recombine. However, in *C. elegans* SC assembly can take place irrespective of homology and consequently a controlling system must be in place to properly coordinate licensing of SC initiation only once homology assessment has been satisfied.

How co-aligning companions recognize each other, remains one of the most important unanswered questions in the field. Although our data do not directly address homology assessment, they have clearly revealed the existence of a novel regulatory pathway that licenses SC initiation once homology is satisfied and releases the constraints to SC polymerization. Furthermore, we show that if homologous synapsis licensing is absent the TZ-like state of meiotic nuclei is sustained. To our knowledge, this is the first mutant condition where a regulatory component is shown to have this phenotype.

We observed that the absence of BRA-2 does not compromise homolog coalignment, as evidenced by largely normal pairing levels (Fig. 2), which were also steadily maintained throughout the pachytene stage, indicating a stable association between the homologs. Synapsis on the other hand, was dramatically impaired, although chromosome ends at the PC were associated with some residual SYP-1 (Fig. 3). This would suggest that while BRA-2 is dispensable to achieve pairing and initial synapsis at chromosome ends, it appears to be essential to transmit a downstream “pairing OK” signal which allows polymerization of the SC along the entire chromosome length.

Previous work has shown that indeed the SC is nucleated at PC sites first and then is polymerized along the entire length of the chromosomes with high processivity^9^.

In line with this, our analysis of GFP::SYP-3 dynamics in live animals has revealed that the formation of SYP-3 nucleation sites is slightly reduced under BRA-2 depletion, however 50% of these foci never undergo elongation. Those that manage to linearize, do so with a strongly reduced elongation rate (Supp. Fig. 2I-L), possibly accounting for the destabilization of the central elements that we observed in BRA-2 depleted animals (Supp. Fig. 2D-G).

The conditional protein degradation allowed by the degron system, combined with the well characterized kinetics underlying nuclear migration within the gonad, allowed us to show that meiocytes entering meiosis without BRA-2 were unable to efficiently assemble the SC, unlike cells that had already entered the pachytene stage in which we observed the robust presence of SYP-1 on aligned chromosomes (Fig. 3). This clearly suggests that BRA-2 promotes SC licensing rather than its stabilization, since in the nuclei undergoing chromosome homology searches without BRA-2 (TZ), synapsis was largely abrogated, while in cells where their homology had been already validated and stabilized by the SC (beyond TZ), loss of BRA-2 did not impede synapsis (Fig. 3). Previous work has shown that pairing takes place independently of the SC at early stages, while becoming essential for stabilizing the association between the homologs later on^40,48,63,64^. The fact that we observed steady state pairing levels in the *bra-2* mutants despite the dramatic impairment in synapsis, further reinforces the dispensable roles exerted by BRA-2 in promoting the initial stabilization of the PCs and emphasizes that its activity is required downstream homologs recognition.

### Chromosome clustering can be sustained upon pairing impairment

Several studies have shown that in *C. elegans*, pairing-defective mutants are often characterized by aberrant/impaired chromosome movement during meiotic prophase. This holds true both in the case of mutations in factors directly involved in the assembly of the SUN/KASH protein bridge, essential to achieve chromosome movement, as well as in motor protein mutants such as the dynein-dynactin family^7,11–13,65^, that transmit the cytoskeletal forces from the microtubules to the SUN/KASH module. Abrogated pairing has also been observed, amongst the others, in mutants of genes involved in regulatory activities of meiotic progression such as *prom-1* and *him-19* mutants^58,59^, in the meiotic kinases mutants CHK-2 and PLK-2^17,20,49^, and in the HORMA-domain containing protein family^23,25,26^. A common denominator in most pairing-defective mutants is represented by lack of a defined region of the gonad containing nuclei with clustered chromatin and deregulated establishment of synapsis, which occurs between non-homologous chromosomes. Lack of synapsis alone on the other hand, triggers prolonged nuclear clustering coupled with extended recruitment of PLK-2 at the PCs and phosphorylation of SUN-1 (a marker for the time window when chromosome movements take place).

Impaired SC formation, however, is not always sufficient to trigger asymmetric chromatin reorganization, as indicated by lack of a defined TZ in the axial element mutants *htp-3* or *him-3,* both structurally essential for the loading of SC central elements^23–25^. This suggests that multiple levels of control are integrated both at the nuclear envelope, as well as at the interface with the chromosome axes to allow chromatin redistribution and prolonged permanence into a homology search state when synapsis and/or recombination are defective.

Amongst axial components, one particular case is provided by the *htp-1* mutants, in which both pairing and chromosome clustering are abrogated as in *htp-3* and *him-3* mutants, however polymerization of the SC takes place between non homologous chromosomes^26^.

Our data show that lack of BRA-2 can bypass the constraints imposed by HTP-1 in acquiring chromatin polarization, but not for achieving chromosome pairing and proper SC assembly, providing as the first unique example of an instance whereby impaired pairing and non-homologous synapsis do not occur as consequential phenomena. The residual synapsis still present upon BRA-2 depletion is further compromised by simultaneous removal of HTP-1, highlighting cooperative roles during SC formation. Therefore, our findings point to the existence of a distinct, HTP-1-independent mechanism that delays exit from TZ when BRA-2 is also absent. Furthermore, these results also suggest that lack of synapsis *per se* may not be the direct trigger sustaining this response, since previous observations reported that neither chromosome clustering nor pairing were restored in synapsis abrogated *htp-1; syp-2* double mutants^26^. This suggests that BRA-2-mediated regulation upon exit from homology search and establishment of synapsis results from an integrated control of multiple signals, including pairing assessment, which in turn harmoniously coordinate early meiotic progression in a timely and faithful manner.

### HIM-17 functions beyond DSB induction

The synthetic effect on SC assembly that we observed under simultaneous removal of BRA-2 and HIM-17 (Fig. 6) was most certainly unexpected. HIM-17 has been identified as one of the SPO11 accessory factors required to achieve efficient DSB formation^33^, and despite more pleiotropic functions having been identified for this chromatin-associated protein in recent years (including regulation of mitotic proliferation versus meiotic entry^66^, transcriptional control of co-opted promoters^67^, positioning of recombination events^68^), no involvement in regulating SC dynamics has so far been reported.

The roles of HIM-17 within the SC pathway are evidently cryptic, since the single mutant does not display any detectable aberrations in pairing or synapsis^33^ (this study), suggesting that its contribution can be appreciated only when synapsis is compromised. This is consistent with both instances in which we found synthetic effects elicited by HIM-17 removal (upon BRA-2 or HTP-1 removal, Fig. 6 and Fig. 7), although the phenotypes that we observed in each case were different, indicating genetic interaction(s) between multiple pathways.

*him-17* mutants depleted for BRA-2 display a complete loss of synapsis, which strikingly impacted on the autosomes but not on the chromosome X, whose pairing and synapsis were instead unaffected. FISH analysis revealed a severe abrogation of pairing of the autosomes, which suggests that the complete loss of synapsis observed in BRA-2-depleted *him-17* mutants may stem from the inability of the homologs to come into proximity, thus preventing SC assembly. This finding is of crucial importance, since it indicates that HIM-17 promotes homolog co-alignment when BRA-2 is absent and that loss of pairing under BRA-2 depletion does not result in non-homologous synapsis. The extensive and extended chromosome clustering observed in BRA-2 and HIM-17 depleted animals, coupled with PLK-2/pSUN-1^S8^ appearance at the nuclear periphery, suggests that the chromosome movement-promoting machinery is in place and that homologs are engaged in an active search for each other.

On the other hand, in *htp-1; him-17* doubles we found only a delay in SYP-1 loading but a strikingly robust and stable recovery of pairing. In this case, it is possible that absence of HIM-17 may just slow down early meiotic progression sufficiently to prevent precocious SC assembly and give the necessary time for the homologs to attain significant levels of co-alignment, which is then stabilized through BRA-2 activity.

This model is consistent with the phenotypes that we observe in the *htp-1* mutants depleted for both BRA-2 and HIM-17 (Fig. 7), which in fact phenocopied BRA-2-HIM-17 depleted animals, both in terms of synapsis and pairing, irrespectively of the presence of HTP-1. This functional crosstalk between BRA-2 and HIM-17 could also be supported by the fact that they establish a physical interaction *in vivo*.

In conclusion, our work provides the first evidence of a complex, multilayered regulatory network tasked with unleashing synapsis to stabilize homolog interactions just at the right time, emphasizing that the signaling that triggers and maintains chromosomes in a clustered and mobile configuration must be protected from the overpowering forces imposed by SC formation.

## MATERIALS AND METHODS

### Strain maintenance and genetics

All strains were cultured according to standard procedures^69^. Worms were grown on nematode growth medium (NGM) plates seeded with OP50 *E. coli* as a food source and maintained at 20°C for all the experiments. To assess embryonic lethality and male progeny, L4 worms of the relevant genotype were individually picked, and mothers were moved onto fresh plates every 24 hours. Dead-eggs/hatched-eggs were scored the day after the mothers were moved and male progeny three days later. CRISPR-tagged strains were generated by SunyBiotech (https://www.sunybiotech.com/), the *him-17::3XHA* tagged line had been previously generated by CRISPR^39^ and the *him-17::GFP* line was generated by biolistic bombardment^33^. A complete list of strains used in this study is reported in Supplementary Table 2.

### Immunofluorescence studies

Worms of the indicated age and genotype were dissected in 15μl of 1XPBS containing 0.1% Tween (PBST) on a 22×22 coverslip. Gonads were fixed by adding 15μl of 2% PFA in 1XPBST and fixation was carried out for 5 minutes at room temperature. Coverslips were flicked off after briefly dipping the slides into liquid nitrogen and samples were transferred to methanol at −20°C for at least 5 minutes. Slides were washed three times in 1X PBST and blocked for 1h at room temperature in 1% BSA in 1XPBST. Primary antibodies were allowed to incubate overnight at room temperature in a humid chamber and the following day, slides were washed three times in 1XPBST for 10 minutes each. Incubation of secondary antibodies was carried out for 2h at room temperature in a humid chamber in the dark. Slides were washed three times in 1XPBST for 10 minutes each in the dark and a 60μl of a DAPI solution in water (2μg/ml) was added on top of the dissected worms. After 1 minute, slides were washed in 1XPBST for at least 20 minutes in the dark, mounted with Vectashield and coverslip were sealed with nail polish. A full list of the antibodies used in this study is reported in Supplementary Table 3.

Samples were acquired with a fully motorized widefield upright microscope Zeiss AxioImager.Z2, equipped with a monochromatic camera Hamamatsu ORCA Fusion, sCMOS sensor, 2304 x 2304 pixels, 6.5 x 6.5 μm size. Z-stacks were set at 0.25μm thickness and images were deconvolved with ZEN Blue Software using the “constrained iterative” algorithm set at maximum strength. Whole projections of deconvolved images were generated with Fiji and processed in Photoshop, where some false coloring was applied.

### Auxin Treatment

Worms of the indicated genotype were selected as L4s or young adults (24h post L4 stage) and picked onto NGM plates with and without 1 mM indole-3-acetic acid (auxin, Sigma). Exposure to auxin was carried out for the indicated times. Given that auxin inhibits bacterial growth, OP50 liquid cultures were concentrated 2X and spotted onto auxin plates a few days in advance and air-dried before placing the worms. Seeded auxin plates were kept refrigerated at 4°C protected from light and used within three weeks.

### Biochemistry

For whole cell extracts, 100 synchronized young adult worms (24h post-L4 stage) were picked into 30 μl of 1XTE (Tris-EDTA Buffer, pH=8) containing 1X complete protease inhibitor cocktail (Roche) and flash-frozen in liquid nitrogen. After thawing, 10 μl of 4X Laemmli Buffer was added, and worms were boiled for 10 minutes with mild shaking. Samples were spun down at maximum speed for 1 minute and run on precast 4-20% gradient acrylamide gels (BioRad) in 1X Tris-Glycine buffer containing 0.1% SDS. Proteins were transferred onto a nitrocellulose membrane (Amersham) in 1X Tris-Glycine buffer containing 20% methanol for 1h at 4°C. Membrane was rinsed in 1XTBS containing 0.1% Tween (TBST) and blocked in 1XTBST containing 5% milk. Incubation of primary and secondary antibodies was carried out in the same buffer overnight at 4°C and 2h at room temperature respectively. For nuclear protein fractionation, extracts were produced as in^70^ without modifications and co-immunoprecipitation was performed by incubating 1-2 mg of nuclear extract with pre-equilibrated anti-HA Affinity Matrix (50 μl, Sigma) or agarose GFP traps (30 μl, Chromotek) in Buffer D (20 mM HEPES pH 7.9, 150 mM KCl, 20% glycerol, 0.2 mM EDTA, 0.2% Triton X-100 and complete Roche inhibitor) overnight at 4°C. The following day, beads were extensively washed in Buffer D and resuspended in 40 μl of 2X Laemmli Buffer and boiled for 10 minutes. Samples were spun down for 1 minute at maximum speed to pellet the beads, and proteins were loaded onto precast 4-20% gradient acrylamide gels.

### In-gel digestion and LC-MS/MS analysis

Selected 1D gel bands were excised manually and after de-staining and washing procedures each band was subjected to protein reduction (10mM DTT in 25mM NH_4_HCO_3_, 45 min, 56°C, 750 rpm) and alkylation (55mM IAA in 25mM NH_4_HCO_3_; 30 min, laboratory temperature, 750 rpm) step. After further washing by 50% ACN/NH_4_HCO_3_ and pure ACN, the gel pieces were incubated with 125 ng trypsin (sequencing grade; Promega) in 50mM NH_4_HCO_3_. The digestion was performed overnight at 37 °C on a Thermomixer (750 rpm; Eppendorf). Tryptic peptides were extracted into LC-MS vials by 2.5% formic acid (FA) in 50% ACN with addition of polyethylene glycol (final concentration 0.001%)^71^ and concentrated in a SpeedVac concentrator (Thermo Fisher Scientific). LC-MS/MS analyses on HIM-17::3XHA pull downs were performed using RSLCnano system (Thermo Fisher Scientific) connected to Orbitrap Fusion Lumos Tribrid (Thermo Fisher Scientific). Prior to LC separation, tryptic digests were online concentrated and desalted using trapping column (300 μm × 5 mm, μPrecolumn, 5μm particles, Acclaim PepMap 100 C18, Thermo Fisher Scientific). After washing of trapping column with 0.1% FA, the peptides were eluted in backflush mode (flow 500 nl/min) from the trapping column onto Acclaim PepMap100 C18 column (3 µm particles, 75 μm × 500 mm; Thermo Fisher Scientific) by 120 min gradient program (mobile phase A: 0.1% FA in water; mobile phase B: 0.1% FA in 80% ACN). Both columns were heated to 40°C. The analysis of the mass spectrometric RAW data files was carried out using the MaxQuant software (version 1.6.10.43) using default settings unless otherwise noted. MS/MS ion searches were done against modified cRAP database (based on http://www.thegpm.org/crap, 112 protein sequences), and UniProtKB protein database for *Caenorhabditis_elegans* (27,028 protein sequences). Trypsin/P enzyme specificity with 2 allowed miss cleavages and minimal peptide length 6 amino acids were set. Only peptides and proteins with FDR threshold <0.01 and proteins having at least one unique or razor peptide were considered.

### EdU staining

Synchronized young adult worms were hand-picked in 50 μl of M9 containing 0.1% tween, to which 50 μl of 1 mM EdU dissolved in DMSO were added before placing the samples at 20°C protected from light.

For the experiment shown in Supp. Fig. 2B, worms were left soaking in M9 containing EdU for 40 minutes, after which they were allowed to recover for 1.5h at 20°C on seeded plates before dissections and staining.

To detect EdU, worms were dissected in 1XPBST and fixed in PFA at 4% final concentration for 10 minutes at room temperature. Coverslips were freeze-cracked in liquid nitrogen and slides were placed in methanol at −20°C for 10 minutes. After three washes in 1XPBST, EdU was detected with the Click-IT reaction kit (ThermoFisher) according to manufacturer protocol.

For the experiment in Fig. 3, worms were incubated in M9 containing EdU for 20 minutes and then placed either on non-auxin plates (untreated and recovery) or auxin plates for the indicated times.

EdU detection combined to immunofluorescence, samples were first processed as for normal immunostaining including incubation with secondary antibodies, after which, the Click-IT protocol was applied according to manufacturer instructions. We and others^72^ noticed that the Click-IT protocol significantly lowers antibody detection efficiency, therefore after testing several markers for the chromosome axes, we chose to employ the anti-phosphoHTP-3^S285^ antibody that we previously generated^73^, which localizes identically as pan anti-HTP-3 but displayed better staining quality.

### Quantification of synapsis, PLK-2, COSA-1 and DAPI bodies

To quantify synapsis, germlines of the indicated genotype were divided into six equal regions spanning the mitotic tip to diplotene entry. Nuclei were considered synapsed only if SYP-1 was fully overlapping with HTP-3 staining. At least three gonads for each genotype were used for quantifications, and the number of nuclei is reported in the Supplementary Table 4.

For PLK-2 quantification, nuclear cell rows from transition zone to late pachytene stage were counted and considered as “positive” only if at least 50% of the cells in a given row displayed PLK-2 staining, as previously shown^57^. At least three germlines were used for the quantification.

For DAPI bodies counts, only −1 and −2 diakinesis nuclei were included and their number for each genotype is reported in Supplementary Table 3.

For OLLAS::COSA-1 staining, only the nuclei in the last seven rows before diplotene stage were included in the quantification^39^. The number of cells scored for the quantifications for each genotype is reported in Supplementary Table 3.

### SUN-1::GFP tracking

For the analysis of SUN-1 aggregates in live animals, worms were anaesthetized with M9 buffer containing 10mM Tetramisole, mounted on 2% agarose pads and covered with coverslips, which were sealed with melted petroleum jelly. SUN-1 aggregates marking chromosome end attachments at the nuclear periphery were filmed as previously described^12^. Before filming adult hermaphrodites, preselected *HA::AID::bra-2; TIR1::mRuby* L4s were incubated at 20°C for 24 hours on plates containing auxin (1mM) or ethanol plates which served as a control. Images were acquired as 0.8µm thick optical sections every 5 seconds for 5 minutes with a DeltaVision Ultra Epifluorescence Microscope, using a 60x silicone objective. The softWoRx suite was used for deconvolution. The images were analyzed using ImageJ (NIH) with StackReg and Manual Tracking plugins.

### GFP::SYP-3 live imaging

Preselected L4s *syp-3(ok758); GFP::syp-3; HA::AID::bra-2; TIR1::mRuby* worms were placed either on an OP50 spread NGM plate or on an NGM plate containing auxin with 3 drops of OP50. The worms were allowed to grow for 24 hours to reach adulthood and begin oogenesis. Live worm mount slides were prepared by placing a 1mm thick 10% agar pad on frosted slides and allowed to cure. At 24 hours post L4 a single one-day old adult was placed in a 0.75µl drop of beads on the agar pad. A cover slip was placed over the pad and the position of the worm marked on the back of the slide. Imaging was done with the DeltaVision deconvolution microscope. A 1024×1024 time lapse 3D stacked image was taken of the full thickness of meiotic nuclei using FITC for 5 minutes. The region imaged was centered on the transition from partial accumulation (foci) of SYP-3::GFP and elongated SYP-3::GFP. After imaging, movies were deconvolved and evaluated for SYP-3::GFP foci initiating an elongation events.

### Fluorescence *In Situ* Hybridization

FISH experiments to detect chromosomes III and V was performed as in^70^ without modifications. To label the pairing center end of chromosome III, the T17A3 cosmid was used to generate a digoxigenin-labelled DNA probe with the Digoxigenin-nick translation mix (Roche) according with manufacturer instructions. The chromosome V was visualized with a biotin-labelled probe generated by amplifying the endogenous 5s rDNA locus by PCR and labelling was performed with the Biotin-nick translation mix (Roche) according with manufacturer instructions. Labelled probes were added to FISH-mix (10% dextran sulfate, 50% formamide, 2XSSCT) and slides were incubated for 3’ at 92°C followed by overnight incubation at 37°C. To combine FISH with SYP-1 staining, we first performed FISH protocol and after probe incubation and post-hybridization washes, slides were washed thrice with 2XSSCT for 10 minutes each at room temperature and blocked in 1% BSA in 2XSSCT for 30 minutes. Incubation with primary chicken anti-SYP-1 antibody^57^ was carried out overnight at room temperature in a humid chamber, and the following day the slides were washed three times with 2XSCCT for 10 minutes each. Goat anti-chicken secondary antibody was combined with FITC-conjugated anti Biotin or Rhodamine-conjugated anti Digoxigenin and left to incubate for 3h at room temperature in the dark. Slides were washed and DAPI staining was performed as for regular immunostaining.

For detection of non-PC of chromosome IV, we employed the same two Cy3-conjugated oligoprobes corresponding to “IV-3” in^74^ and performed FISH “in tube” with minor modifications. Briefly, worms were dissected on a 22×22 coverslip in 30 μl of 1XEGG buffer containing 0.1% Tween and fixed by adding an equal volume of 8% PFA in 1XEGG buffer containing 0.1% Tween. Samples were fixed for 4 minutes at room temperature and then transferred to 1.5-mL Eppendorff tubes, in which 1 mL of cold methanol was added. Tubes were placed at −20°C until completion of dissections. Methanol was replaced with 1 mL of 2XSSC containing 0.1% Tween to perform three washes at room temperature. After the last wash, samples were transferred to 0.1 mL PCR tubes and 100 μl of 50% formamide in 2XSSC containing 0.1% Tween was added to carry out a pre-hybridization step of the samples in a PCR thermocycler at 37°C overnight. The following day, formamide was removed and 40 μl of FISH mix containing 2 μl of each oligoprobe from a 10 μM stock were added. Probe hybridization was carried out in a thermocycler at 92°C for 2’30’’, 72°C for 2’ and 37°C overnight. The following day, FISH mix was removed, samples were washed three times in 2XSSCT for 5 minutes each at room temperature in the dark. DAPI was allowed to stain in the tubes for 2 minutes and after two 10-minutes wash in 2XSCCT, the dissected germ lines were transferred to a superfrost positively charged slide in minimal volume and mounted with 12 μl of Vecatshield.

## ACKNOWLEDGMENTS

We are grateful to E. Martinez-Perez, S. Arur, A. Villeneuve, W. Kelly, Y. Kim, E. Kipreos, M. Colaiacovo and A. Dernburg for valuable strains and reagents. We thank Chantal Wicky for sharing unpublished data and Judith Yanowitz for critical discussions. We are grateful to Anne Villeneuve for critical reading of the manuscript. We acknowledge the core facility CELLIM supported by the Czech-BioImaging large RI project (LM2023050 funded by MEYS CR) for their support with obtaining scientific data presented in this paper. We acknowledge CIISB, Instruct-CZ Centre of Instruct-ERIC EU consortium, funded by MEYS CR infrastructure project LM2023042, for the financial support of the measurements at the CEITEC Proteomics Core Facility. Computational resources were provided by the e-INFRA CZ project (ID:90254), supported by MEYS CR. *In vivo* filming of chromosome-nuclear envelope end attachments was performed by the BioOptics Facility at Max Perutz Labs using the VBCF instrument pool. Some strains were provided by the CGC, which is funded by NIH Office of Research Infrastructure Programs (P40 OD010440). Research in the Silva lab is funded by the Czech Science Foundation (GA23-04918S), the Smolikove lab is funded by NSF (2027955), the Jantsch lab is funded by the FWF (Fonds zur Förderung der wissenschaftlichen Forschung) [SFB F 8805-B] and the Zetka lab by the Canadian Institutes of Health Research (PJT-173381).

## Supplementary Figures Legends

**Supp. Figure 1. (A)** Top: Western blot on whole-cell protein extracts showing HA::AID::BRA-2 depletion efficiency upon exposure to auxin at the indicated times. Actin was used as loading control. Bottom: quantification of protein depletion from two biological duplicates. Actin was used for normalization. **(B)** HTP-3 and SYP-1 staining in control animals compared to *bra-1(nk1)*, *bra-2(ok1171)* and *bra-2(ok1171); bra-1(nk1)* double mutants, showing that BRA-1 does not contribute to establishment of synapsis. Scale bar 10 μm.

**Supp. Figure 2. Loss of BRA-2 destabilizes SC central elements. (A)** Co-staining of mitotic markers WAPL-1/CYE-1 shows normal mitosis-meiosis switch in control animals and *bra-2(ok1171)* mutants. Scale bar 5 μm. **(B)** Left: representative images of EdU staining in the nuclei of the mitotic distal tip of the germ line counterstained by DAPI. Scale bar 5 μm. Right: quantification of the fraction of EdU-positive cells over the total number of nuclei shows that absence of BRA-2 does not impair mitotic replication capacity. Bars indicate SD and statistical significance assessed by the T test (*ns*= non-significant). **(C)** Representative images of TZ-EP nuclei stained for HTP-3/SYP-1 in *HA::AID::bra-2; TIR1::mRuby* animals before and after auxin exposure, showing identical SC-impaired phenotype as *bra-2(ok1171)* deletion mutants. Scale bar 10 μm. **(D-F)** Early pachytene nuclei from the indicated genetic backgrounds showing that removal of BRA-2 upon auxin exposure impairs loading of SYP-2, SYP-3 and SYP-4. Scale bar 5 μm. **(G)** Western Blot analysis on whole-cell extracts indicates destabilization of SYP-2/-4 upon BRA-2 depletion. Actin was used as loading control. **(H)** Quantitative analysis of *syp-1* and *syp-4* transcripts by qPCR shows normal levels in both WT and *bra-2(ok1171)* mutant worms. **(I)** *In vivo* imaging of GFP::SYP-3 in untreated worms and after exposure to auxin indicates reduced formation of SYP-3 nucleation sites, **(J)** impaired elongation proficiency and **(K-L)** starkly reduced SYP-3 polymerization speed. Statistical significance was calculated by the Mann-Whitney test.

**Supp. Figure 3. BRA-2 depletion impairs licensing of the SC rather than its maintenance. (A)** SYP-1/HTP-3 immunostaining on untreated worms and upon prolonged BRA-2 depletion (48h) shows SC defects throughout the germ line. The yellow square depicts nuclei in transition zone/early pachytene, blue square indicates nuclei at the mid-late pachytene stage. Scale bar 10 μm. **(B)** Western blot on whole-cell extracts indicates that upon exposure to auxin, levels of HA::AID::BRA-2 are fully recovered after 72h. Actin was used as loading control.

**Supp. Figure 4.** Loss of BRA-2 suppresses precocious exit from early meiotic stages in *htp-1* mutants. Top: whole-mount gonads stained for PLK-2 and counterstained by DAPI. BRA-2 depletion in *htp-1* nulls restores retention of PLK-2 at the nuclear periphery and extended nuclear clustering, indicating delayed exit from transition zone. Scale bar 10 μm.

**Supp. Figure 5. BRA-2 is enriched on the autosomes and does not engage into epistatic relationships with BRA-1. (A)** Co-staining of HA::AID::BRA-2 with different autosome-associated factors. Scale bar 5 μm. **(B)** Localization of BRA-2 is independent of DSBs and synapsis. Scale bar 5 μm. **(C)** Top: whole-mount gonad showing distinct temporal expression patterns of HA::AID::BRA-2/FLAG::BRA-1. Bottom: localization of BRA-1/BRA-2 is not interdependent. **(D-E)** Western blot on whole-cell extracts showing that BRA-1-BRA-2 stability is not interdependent.

**Supp. Figure 6. HIM-17 promotes synapsis in absence of BRA-2. (A)** SYP-1/HTP-3 immunostaining shows a dramatic impairment of synapsis in absence of BRA-2 and HIM-17. **(B)** SYP-1/HIM-8 immunostaining showing that only the chromosome X retains synapsis in mid-late pachytene cells. Scale bar 10 μm. **(C)** Immunofluorescence showing mutual exclusive loading of WAPL-1/CYE-1 in the premeiotic region of the gonad and at meiotic entry, indicating that cell cycle progression is not impacted by loss of BRA-2 and HIM-17. Scale bar 10 μm. **(D)** SYP-1/HTP-3 immunostaining reveals that SC phenotype is not worsened in absence of *him-5* or **(E)** *xnd-1* upon BRA-2 depletion. Scale bar 5 μm.

**Supp. Figure 7. Loss of HIM-17 largely restores homologs pairing in *htp-1* mutants. (A)** Representative images of Ch. V FISH in mid-pachytene nuclei in the indicated mutants. Scale bar 5 μm. **(B)** Ch. V FISH and SYP-1 immunostaining in the indicated mutant backgrounds. Scale bar 5 μm. **(C)** Early pachytene cells from *him-17::AID; TIR1::mRuby* tagged animals stained with anti-AID and RAD-51 antibodies indicates normal HIM-17 expression and DSB formation. Scale bar 10 μm. **(D)** Whole-mount gonads stained with anti-pSUN-1^S8^ antibodies and counterstained by DAPI. Impaired pairing observed under BRA-2-HIM-17 double depletion does not depend on defective chromosome movement. Scale bar 10 μm.

