## Supplementary figures and images for "Distinct intersecting pathways link homolog pairing to initiation of meiotic chromosome synapsis"

### Supplemental Figure 1

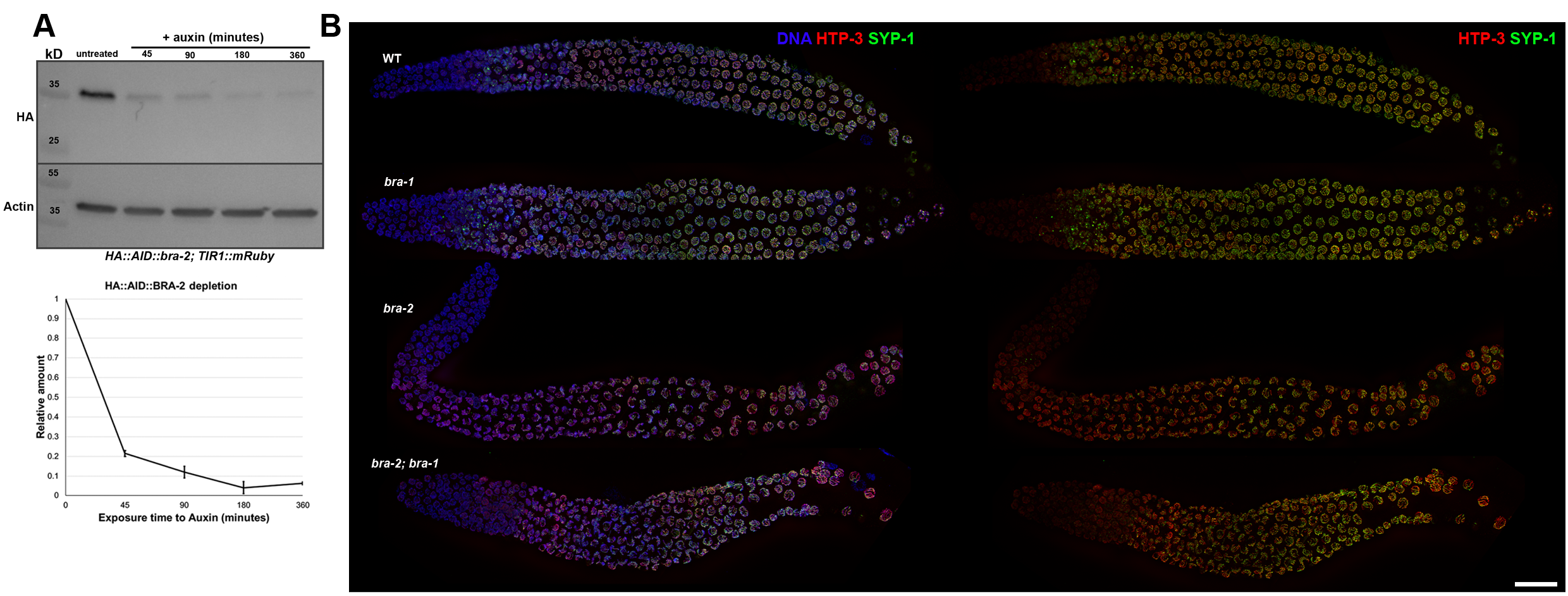

### Supplemental Figure 2

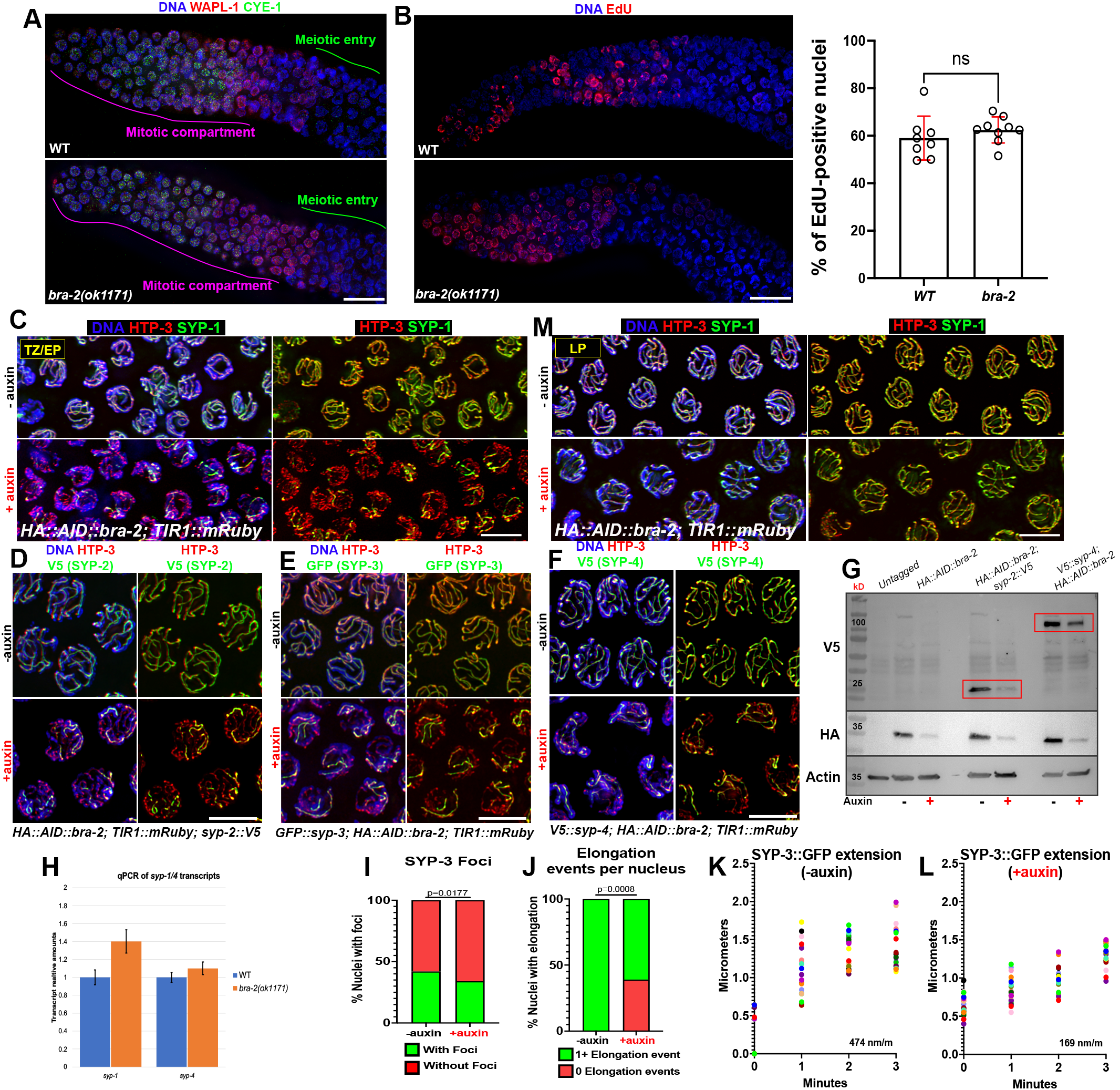

### Supplemental Figure 3

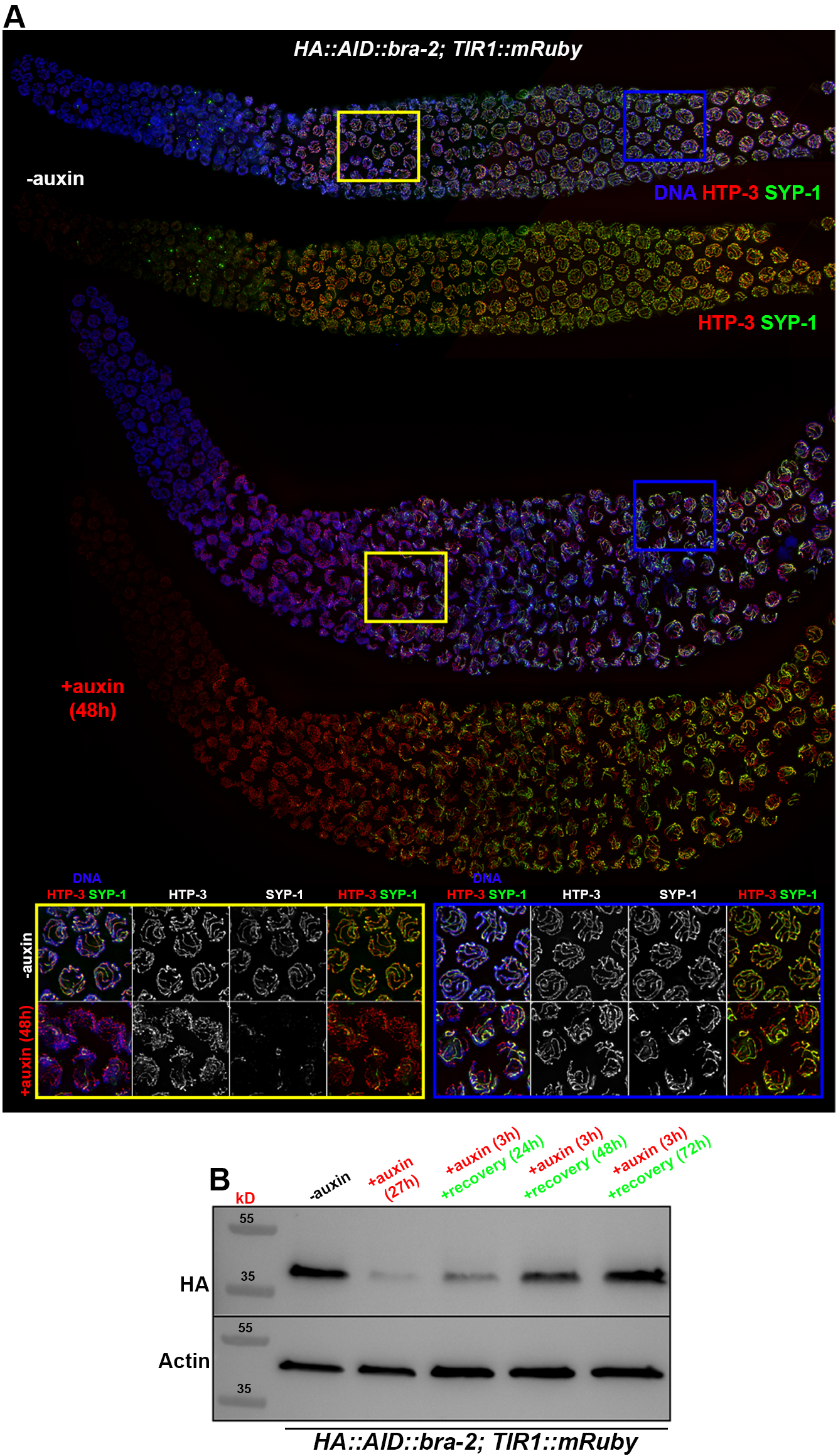

### Supplemental Figure 4

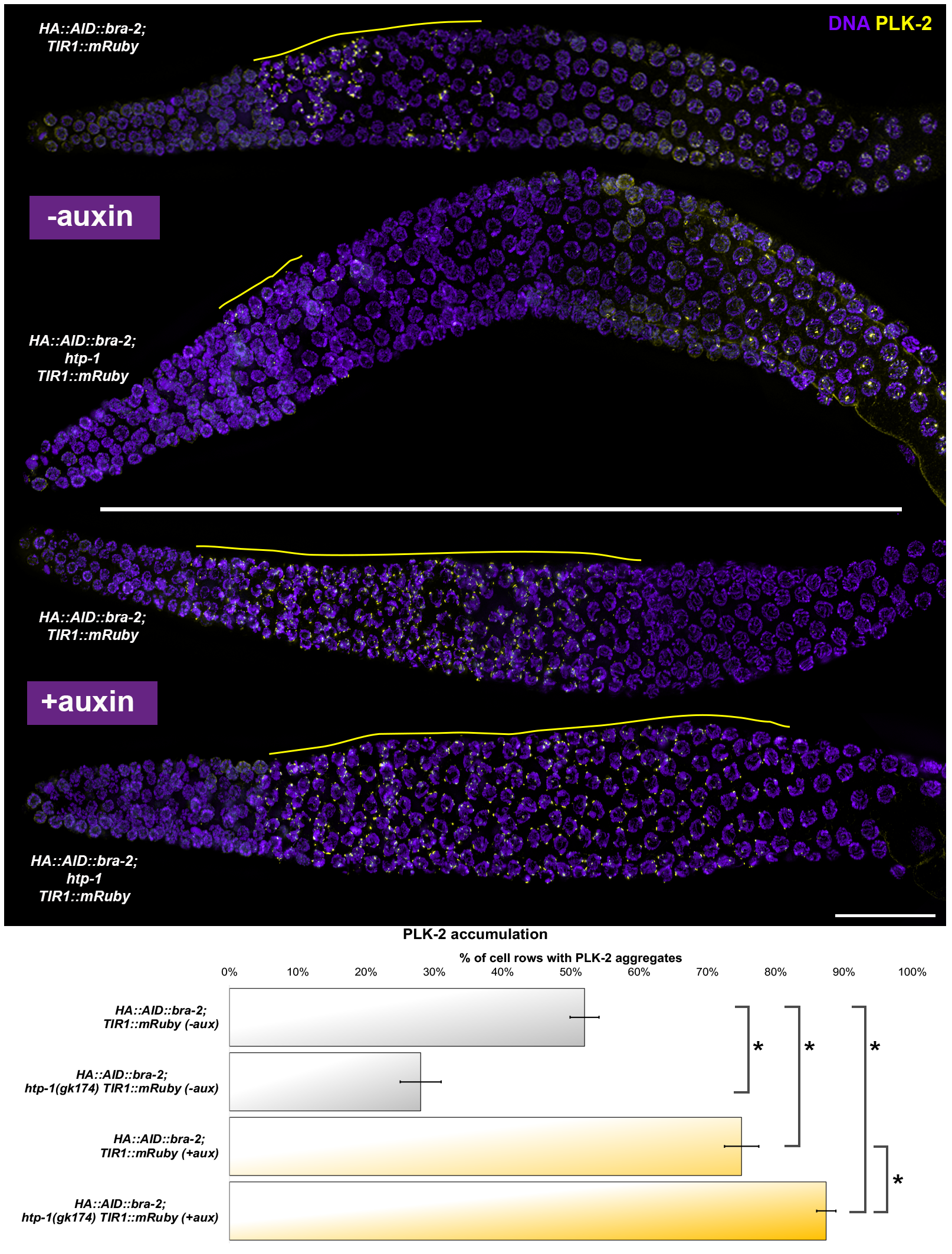

### Supplemental Figure 5

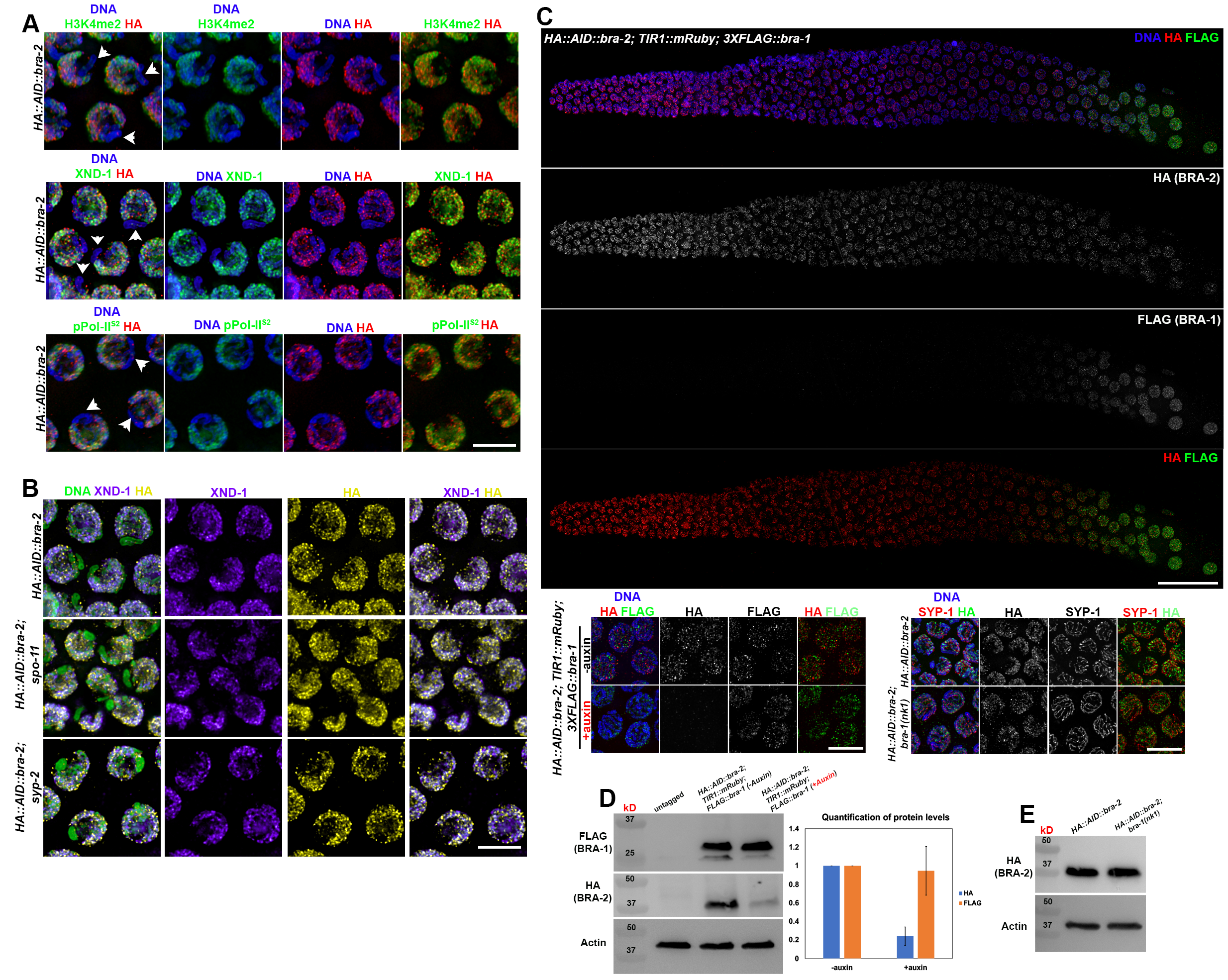

### Supplemental Figure 6

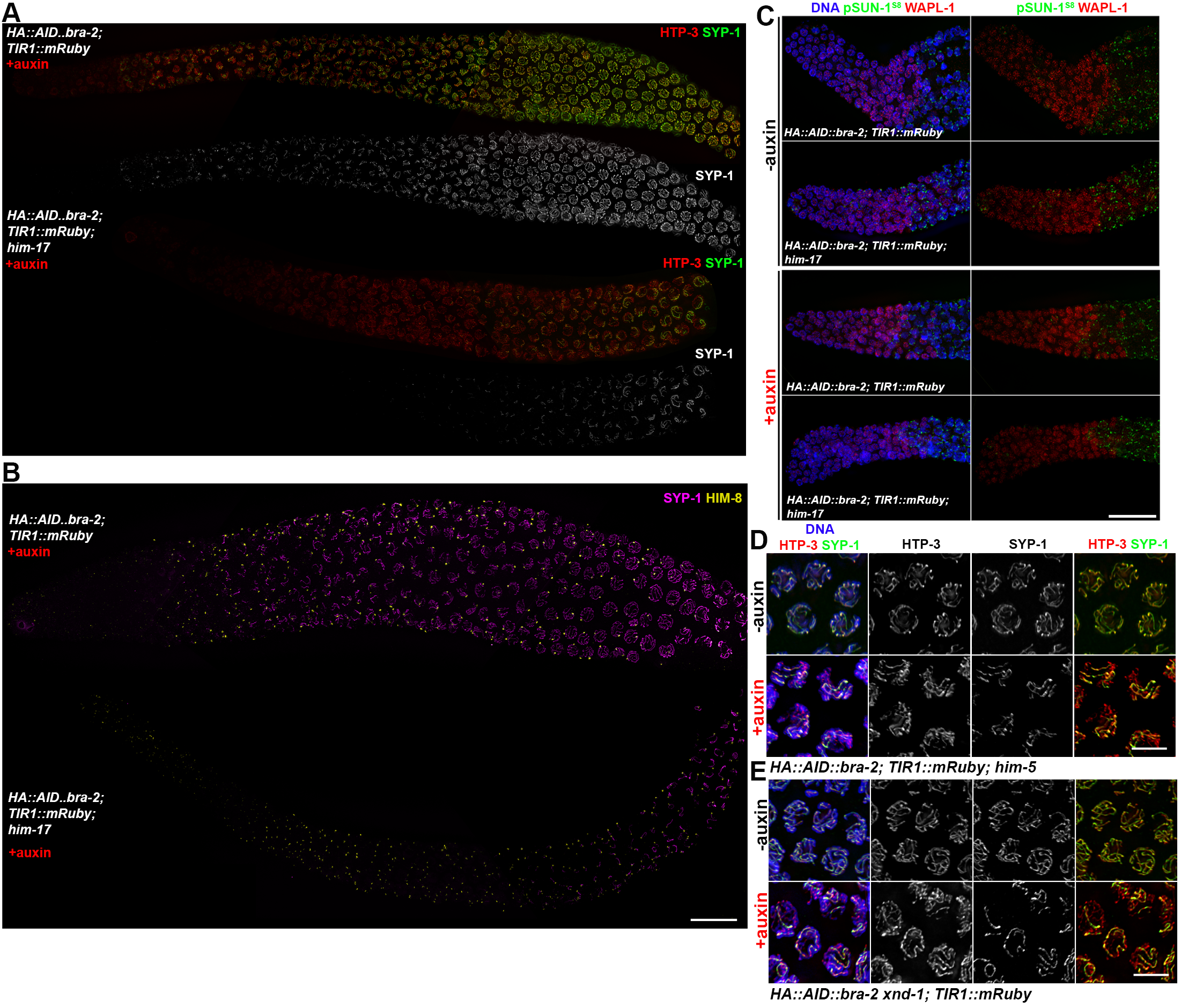

### Supplemental Figure 7

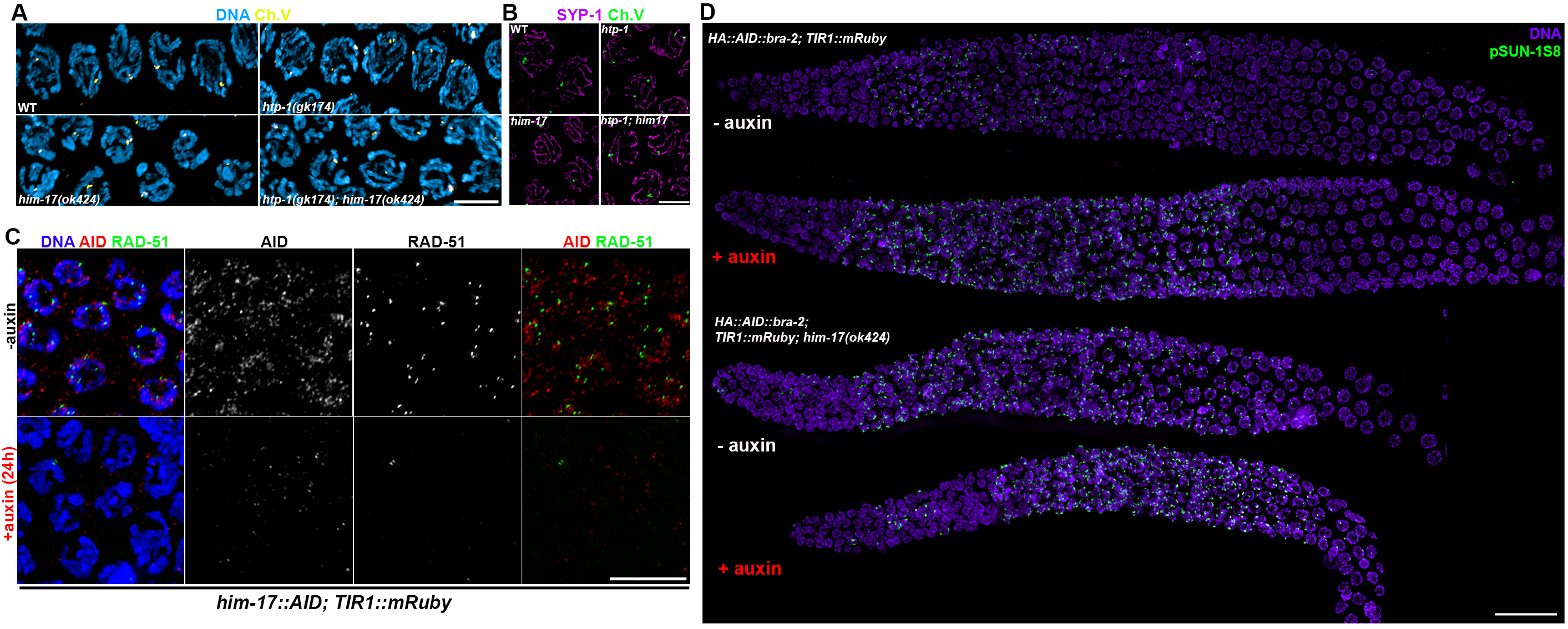
